# Noradrenergic signaling controls Alzheimer’s disease pathology via activation of microglial β2 adrenergic receptors

**DOI:** 10.1101/2023.12.01.569564

**Authors:** L. Le, A.M. Feidler, H. Li, K. Kara-Pabani, C. Lamantia, M.K. O’Banion, K. A Majewska

## Abstract

In Alzheimer’s disease (AD) pathophysiology, plaque and tangle accumulation trigger an inflammatory response that mounts positive feed-back loops between inflammation and protein aggregation, aggravating neurite damage and neuronal death. One of the earliest brain regions to undergo neurodegeneration is the locus coeruleus (LC), the predominant site of norepinephrine (NE) production in the central nervous system (CNS). In animal models of AD, dampening the impact of noradrenergic signaling pathways, either through administration of beta blockers or pharmacological ablation of the LC, heightened neuroinflammation through increased levels of pro-inflammatory mediators. Since microglia are the resident immune cells of the CNS, it is reasonable to postulate that they are responsible for translating the loss of NE tone into exacerbated disease pathology. Recent findings from our lab demonstrated that noradrenergic signaling inhibits microglia dynamics via β2 adrenergic receptors (β2ARs), suggesting a potential anti-inflammatory role for microglial β2AR signaling. Thus, we hypothesize that microglial β2 adrenergic signaling is progressively impaired during AD progression, which leads to the chronic immune vigilant state of microglia that worsens disease pathology. First, we characterized changes in microglial β2AR signaling as a function of amyloid pathology. We found that LC neurons and their projections degenerate early and progressively in the 5xFAD mouse model of AD; accompanied by mild decrease in the levels of norepinephrine and its metabolite normetanephrine. Interestingly, while 5xFAD microglia, especially plaque-associated microglia, significant downregulated *β2AR* gene expression early in amyloid pathology, they did not lose their responsiveness to β2AR stimulation. Most importantly, we demonstrated that specific microglial β2AR deletion worsened disease pathology while chronic β2AR stimulation resulted in attenuation of amyloid pathology and associated neuritic damage, suggesting microglial β2AR might be used as potential therapeutic target to modify AD pathology.

## Introduction

Alzheimer’s disease (AD) is characterized by progressive accumulation of Aβ plaques and neurofibrillary Tau tangles, triggering a chronic neuroinflammatory response that mounts positive feed-back loops between inflammation and protein aggregation, aggravating neurite damage and neuronal death. While AD is commonly associated with cognitive impairments that result from neocortical and hippocampal atrophy, one of the earliest brain regions to undergo neurodegeneration in AD is the locus coeruleus (LC), a small nucleus located in the brainstem responsible for most of the norepinephrine (NE) production in the central nervous system (CNS). Neurons of the LC project to and innervate the cerebral cortex, hippocampus, and cerebellum, where NE is released and diffuses over widespread areas. NE can act on adrenergic receptors (ARs) in both neurons and glia ^1^, regulating a wide array of critical brain functions including cognition, attention, emotion, and the sleep-wake cycle ^2^. While some LC neuronal loss is common in normal aging ^3,4^, the degree of LC degeneration is much more pronounced in AD patients with an estimated increase of 50-80% in neuronal death compared to age-matched controls ^4–7^.

In animal models of AD, lowering NE signaling through either pharmacological ablation of LC neurons ^8–10^ or blocking of NE receptors ^11,12^ augments both neuroinflammation and Aβ plaque load. In contrast, partially rescuing endogenous NE signaling via prolonged exposure to β-AR agonists ^13^ or the NE precursor L-Threo-3,4-dihydroxyphenylserine ^14^ reduces levels of several inflammatory cytokines and improves amyloid pathology. Moreover, pharmacological lesioning of the LC also decreases microglial recruitment to Aβ plaques ^9^, thus reducing microglia’s ability to “wall-off” toxic Aβ deposits and potentially restrict neuritic damage ^15,16^. NE signaling can also directly inhibit microglia reactivity in acute inflammation contexts *in vitro* ^17–19^. Thus, mounting evidence indicates that microglia are responsible for translating the loss of NE tone in AD patients to chronic neuroinflammation and the associated exacerbation of disease pathology.

Although microglia express both α- and β-ARs ^1,17,20^, NE likely exerts its anti-inflammatory effects through the activation of microglial β -ARs. Pharmacological agonism of these receptors reduces lipopolysaccharide (LPS)-induced microglial production of inflammatory cytokines such as TNF-α and IL-6 in slice and primary microglia cultures ^17–19^. Interestingly, in the healthy brain, β2ARs are highly enriched in microglia relative to other types of ARs and, in fact, microglia express β2ARs at levels much higher than that seen in other CNS cell types ^21,22^, suggesting that microglia might have unique responses to NE through these receptors. Indeed, previous work from our lab and that of other labs showed that endogenous NE inhibits microglial surveillance and process motility via microglial β2AR during wakefulness, suggesting a less immune-vigilant functional state ^23,24^. Furthermore, single-cell RNA-sequencing in aged ^25^ and 5xFAD mice ^26^, a commonly used AD animal model, revealed a significant downregulation of β2AR expression in the age-associated microglia and the disease-associated microglia (DAM) clusters, respectively. DAM are localized around Aβ plaques and are transcriptionally defined by downregulation of homeostatic genes, with concomitant upregulation of pro-inflammatory and lysosomal phagocytic genes ^26,27^. These results suggest a potential relationship between the DAM phenotype and β2AR expression, prompting the question of whether there is a spatiotemporal relationship between microglial β2AR expression and amyloid pathogenesis.

Although some studies have shown LC abnormalities in 5xFAD mice ^14^ and progressive degeneration of the LC-NE system in another common amyloidosis model APP/PS1 ^28^, to the best of our knowledge, there has not been a systemic characterization of dynamic changes in both endogenous LC-NE signaling and microglial responsiveness to NE throughout amyloid pathogenesis. In this study, we utilized the 5xFAD mouse model of AD and observed an early degeneration of NE projections followed by LC neuronal loss at more advanced disease stages, accompanied by mild decreases in the levels of norepinephrine and its metabolite normetanephrine in the brain. Interestingly, we found that 5xFAD microglia downregulated their expression of β2AR mRNA early and became insensitive to anesthesia which affects endogenous NE, an effect that was most apparent in plaque-associated microglia. Our studies also illuminated the important role of β2AR signaling specifically within microglia by showing opposing effects of increasing and decreasing this signaling on amyloid pathology, using pharmacological and transgenic approaches. Taken altogether, our results suggested the potential of microglial β2AR as a specific target to leverage for disease modifying therapy.

## Results

The goal of this study was to determine how β2AR signaling, specifically within microglia, affects AD pathology. We chose to focus on amyloid pathology and utilized the commonly used 5xFAD mouse model of amyloidosis. To determine the time course of changes in endogenous NE release and microglial response to NE, we examined three different age groups which correspond to early pathology (4 months old), established amyloid pathology (6 months old) and late phase of disease progression when plaque load plateaus (9 months old) ^29,39^. This allowed us to tie changes in microglial NE signaling to amyloid deposition in a spatiotemporal manner, setting the stage for experiments that manipulated microglial β2AR signaling while visualizing changes in Aβ plaques.

### NE neurons in the LC degenerate late in 5xFAD mice

To characterize changes in endogenous NE signaling as a function of amyloid pathology, we first determined the degree of NE releasing LC neuron loss in 5xFAD mice compared to littermate controls across the 3 different age groups (4, 6 and 9 months) (Fig. 1).

**Fig. 1:**
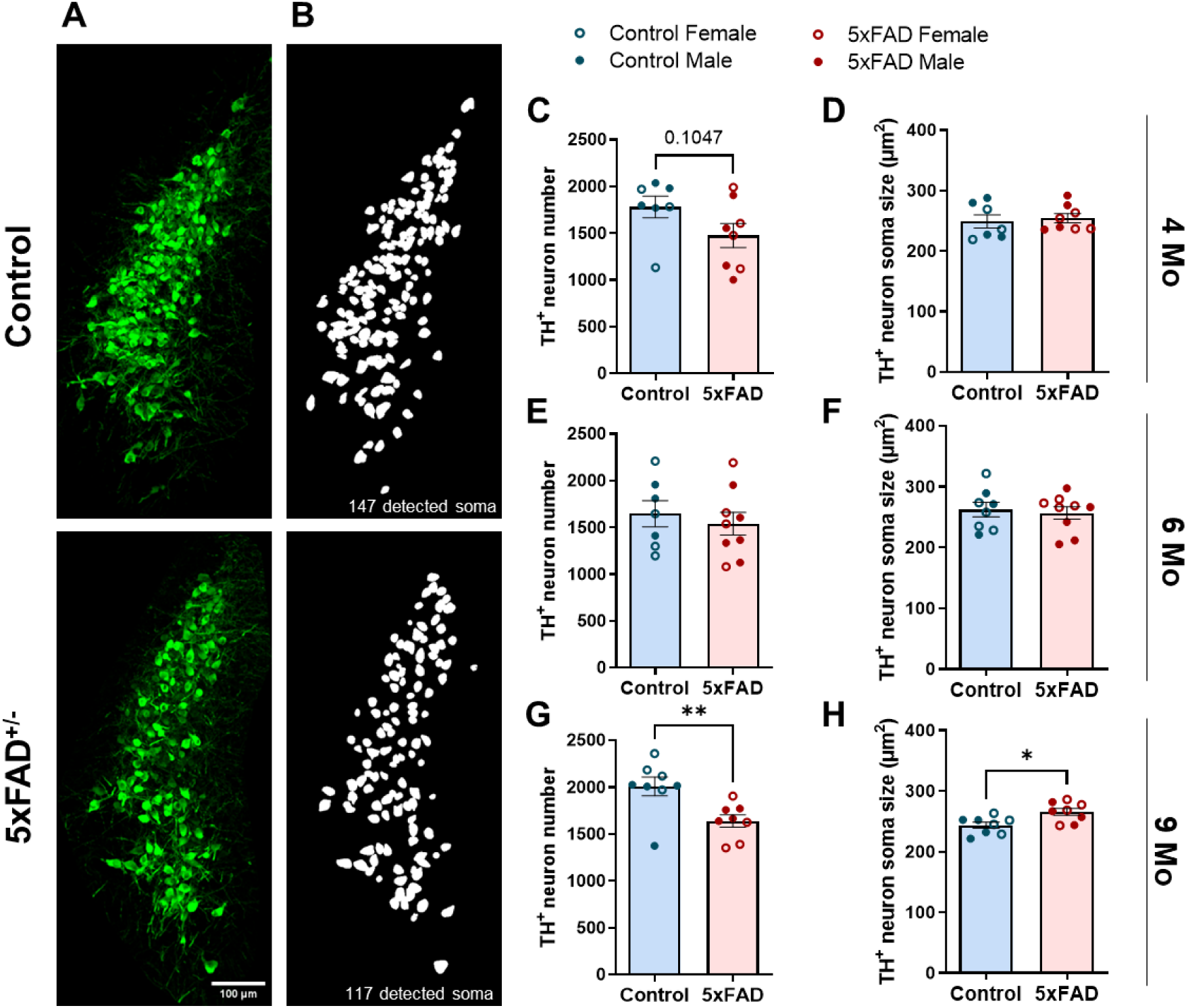
Loss of NE neurons in the LC in 5xFAD mice at advanced amyloid pathology stage. Representative 20x confocal images of LC neurons immunolabeled with TH (**A**). Representative soma detection using Cellpose custom-trained model (**B**). Number and size of LC neurons in 4-month (**C-D**), 6-month (**E-F**), and 9-month (**G-H**) control and 5xFAD mice. *n = 7-9, Student t-test; *p<0.05, **p<0.01. Scale bar: 200μm*.

Immunofluorescent staining for TH^+^ cells was carried out in all brain sections containing the LC, and the total number of TH^+^ cells within the LC was quantified using a custom-trained Cellpose model. This revealed significant LC neuronal loss in 5xFAD mice compared to wild-type (WT) control littermates at 9 months of age (Fig. 1G) alongside a significant increase in soma size of surviving neurons (Fig. 1H). No differences between the two genotypes were detected at 4 or 6 months of age (Fig. 1C-F). Due to the large number of brain slices used for LC quantification, immunohistochemistry experiments were carried out independently for each age group. Thus, variation among age groups most likely reflects the intrinsic variability of immunolabeling and tissue quality among the different experiments. The observed degeneration of LC neurons was not directly attributable to the increase in amyloid pathology at older ages because the LC is relatively devoid of dense core plaques (Supplementary Fig. 1A). However, we detected a significant increase in Iba1 and GFAP immunolabelling in the LC of 6-month-old 5xFAD mice compared to WT littermates, suggesting enhanced neuroinflammation caused by reactive microglia and astrocytes. These elevated levels of reactive glia preceded the reduction in LC neurons in 9-month-old 5xFAD mice, and while higher reactivity was also seen in 9-month-old animals this effect was not statistically significant (Supplementary Fig. 1C-D, F-G).

### Reduced cortical NE levels in 5xFAD mice

To examine whether LC neuron loss translates to a reduction in brain levels of NE, we measured cortical NE concentration in 5xFAD and WT mice in all three age groups using ELISA. Interestingly, we observed an early reduction in cortical NE levels in 5xFAD compared to WT mice at 4 months of age (Supplementary Fig. 2A), despite the fact that LC neuronal loss was not observed until 9 months (Fig. 1G). Furthermore, the levels of cortical NE were similar between 5xFAD and WT at 6 and 9 months of age (Supplementary Fig. 2A). Because NE released in the brain is rapidly degraded by the abundant post-synaptic membrane-bound catechol-O-methyltransferase (COMT) into its more stable metabolite normetanephrine (NMN) ^40,41^, we also measured levels of cortical NMN as a proxy for levels of released NE. Unlike the early reduction in levels of NE, levels of NMN were similar between 5xFAD and WT mice at 4 months; however, we observed a mild but consistent decrease in NMN levels in 5xFAD mice compared to WT in older age groups which did not reach statistical significance (Supplementary Fig. 2B). Together, these findings suggest dysfunction in NE signaling along with compensatory mechanisms that may stabilize the levels of released NE throughout the course of amyloid pathology progression.

### TH^+^ nerve fibers degenerate early in the brains of 5xFAD mice

Given that the number of LC neurons decreases late in amyloid pathology progression, we sought to determine whether the loss of NE projections to the cortex could explain the earlier decreases in cortical NE and NMN. We quantified TH^+^ nerve fibers as putative noradrenergic projections, alongside Aβ plaques and neuroinflammatory markers (Fig. 2A). We focused on the anterior cingulate cortex (ACC) and primary visual cortex (V1), as the former is one of the earliest areas in the brain to exhibit amyloid plaque deposits ^29,39^ while the latter is accessible for *in vivo* assessment of microglia dynamics (see below). In ACC, senile amyloid plaque deposition and expression of markers of microglial and astrocytic reactivity were already high at 4 months of age and only GFAP exhibited further increases with aging (Fig. 2B-D). In V1, plaque deposition grew from 4- to 6-month-old, alongside increases in astrocytic reactivity markers, while microglial reactivity remained increased as compared to WT controls at all ages (Fig. 2F-H). In both brain areas, female 5xFAD mice exhibited more pronounced amyloid pathology than their male counterparts (Fig. 2B, F). It should be noted that here we only quantified senile, dense core plaques with Methoxy-X04 immunofluorescence and did not make a conclusion on the overall amyloid pathology which includes other Aβ species. Previous findings ^29,39^ and our own observation of graded increases in GFAP levels with age indicate overall age-related exacerbation of amyloid pathology in the ACC and V1 in 5xFAD mice. In ACC we observed a significant decrease in TH^+^ projections at 4 months, and in V1 at 6 months in 5xFAD compared to age-matched WT mice (Fig. 2E, I), suggesting that the degeneration of NE projections correlates with the onset of substantial amyloid pathology and neuroinflammation. At 9 months, however, the decrease in TH^+^ nerve fibers in WT mice matched that of 5xFAD mice, possibly suggesting that the process of aging also negatively influences the maintenance of NE fibers irrespective of amyloid pathology.

**Fig. 2:**
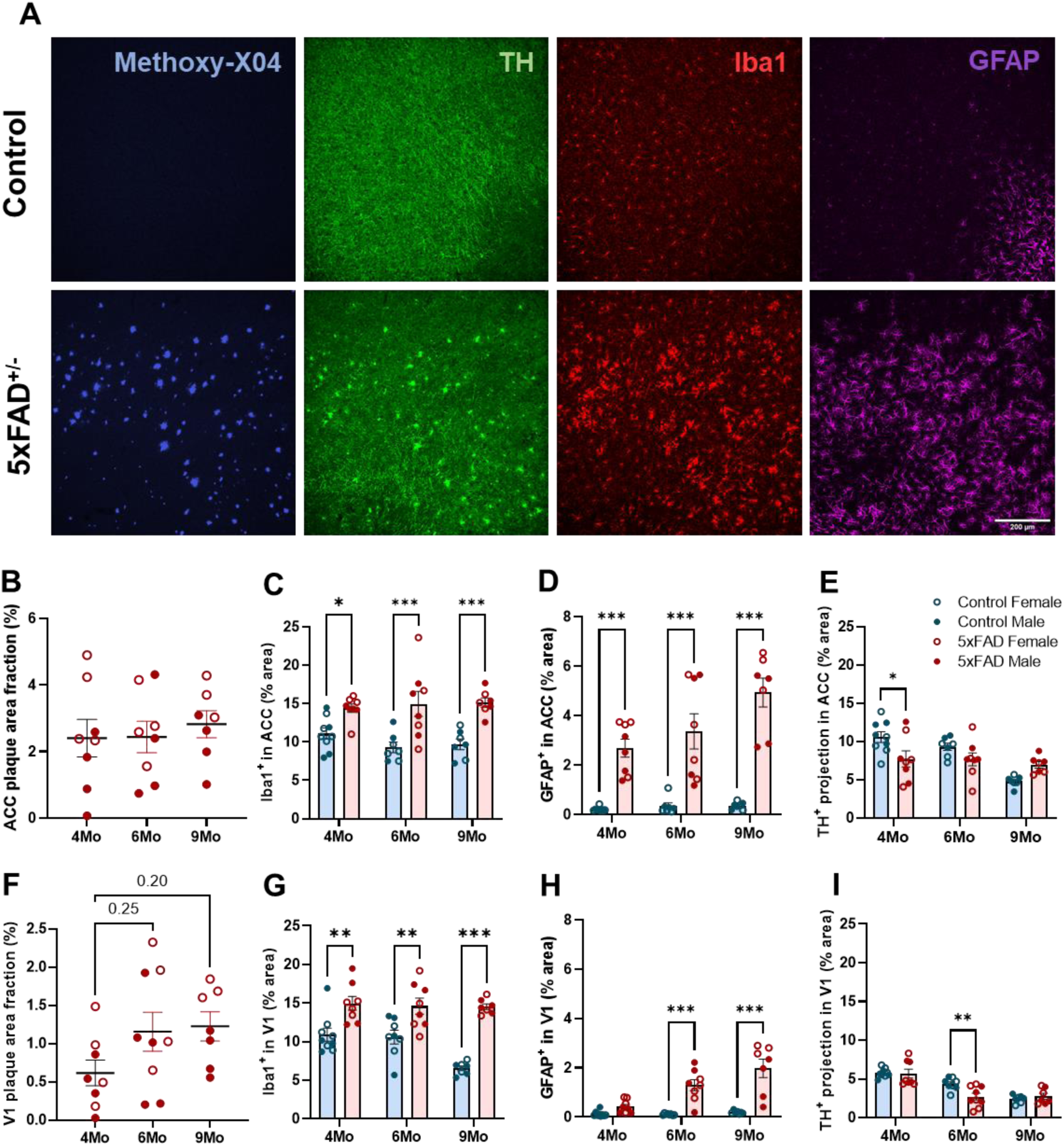
Early loss of TH^+^ projection in cortex in 5xFAD mice. Representative 20x confocal images showing neuritic plaques (Methoxy-X04, blue), NE projections immunolabeled with TH (green), microglia (Iba1, red), and astrocytes (GFAP, magenta) in the ACC (**A**). Analyses were done in ACC (**B-E**) and V1 (**F-I**). Quantification of senile plaque load labeled with Methoxy-X04 (**B, F**). Area fraction of both microglia (**C, G**) and astrocyte (**D, H**) increased early in 5xFAD mice. Decrease in TH^+^ projections in ACC at 4 months (**E**) and in V1 at 6 months (**I**) in 5xFAD mice compared to littermate controls. *n = 7-9, one-way ANOVA (**B, F**) or two-way ANOVA (**C-E, G-I**) with Bonferroni correction; *p<0.05, **p<0.01, ***p<0.001. Scale bar: 200μm*.

### The modulation of microglial surveillance by anesthesia is affected by age and amyloid pathology

NE is released at higher levels in the awake state; while under anesthesia, NE concentrations in the cortex decrease, augmenting microglial dynamics in young adult mice ^23^. Thus, to determine how the loss of endogenous noradrenergic signaling impacts microglia behavior, we imaged microglial dynamics in animals expressing the CX3CR1^GFP^ transgene (CX3^G/+^), which allows for fluorescent labeling of microglia during wakefulness and under anesthesia using a chronic cranial window preparation over V1 (Fig. 3A). Similar to our previous work ^23^, we found that cortical microglial process surveillance significantly increased when a young (4-6 months old) adult CX3^G/+^ animal was anesthetized compared to when it was awake (Fig. 3B-C). Microglia in 5xFAD animals, on the other hand, exhibited different response depending on their proximity to amyloid plaques. Microglia distal from Aβ plaques increased their parenchymal surveillance under anesthesia though to a lesser extent than microglia in age-matched controls, while plaque-associated microglia did not alter their dynamics under anesthesia (Fig. 3B-C). At 9 month of age, microglial surveillance was not affected by anesthesia in CX3^G/+^ or 5xFAD/CX3^G/+^ mice (Fig. 3D). Interestingly, microglial arbors were less ramified in awake compared to anesthetized state across all age groups (Fig. 3G-I).

**Fig. 3:**
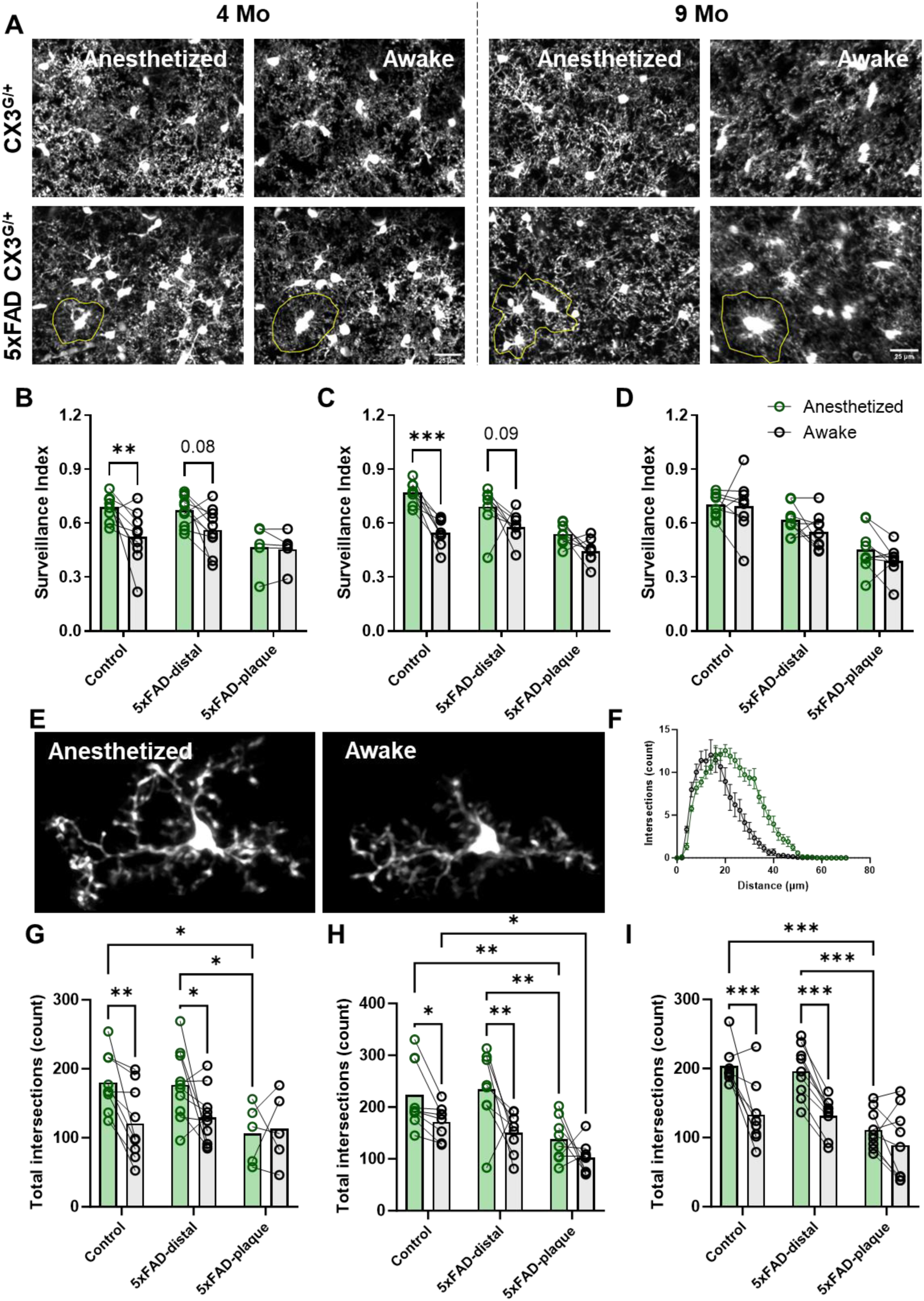
Anesthesia (lower NE) affects microglia surveillance differently with pathology and age in V1. Representative *in vivo* two-photon images showing fewer microglial pixels in time projected images in awake animals (reflecting lower microglial surveillance) than in anesthetized animals at 4-month (left 4 panels) but not as pronounced at 9-month (right 4 panels) of age (**A**). Quantification of microglial surveillance in awake versus anesthetized state in CX3CR1^GFP/+^ (Control) and CX3CR1^GFP/+^ 5xFAD^+/-^ (5xFAD) mice (**B:** 4 months**; C:** 6 months **; D:** 9 months). Representative of manually-selected individual microglia from awake and anesthetized mice for Sholl analysis (**E**). Representative Sholl curves showing Sholl profiles of microglia in awake (black) and anesthetized (green) control mice at 4 months old (**F**). Both control and plaque-distal microglia are more ramified (higher total intersections) under anesthesia (**G:** 4 months**; H:** 6 months **; I:** 9 months). *n = 8-10, repeated measures ANOVA with Bonferroni (B-D) or Tukey (G-I) correction, **p<0.01, ***p<0.001. Scale bar: 25μm*.

### Microglial expression of homeostatic markers and β2AR is downregulated with age and increasing amyloid pathology

The differences in the responses of plaque-associated versus plaque-distal microglia to endogenous NE release in awake 5xFAD mice prompted us to determine whether microglia themselves lose sensitivity to NE signaling as a result of Aβ plaque proximity. In order to separate plaque-associated and plaque-distal microglia, we injected mice with MX04, a brain-permeable fluorescent probe for Aβ, 24 h prior to sacrifice and FACS-sorted cortical CD11b^+^CD45^int^ microglia into MX04^+^ and MX04^−^ fractions (Supplementary Fig. 3; Fig. 4A). In agreement with the previously reported downregulation of homeostatic markers in 5xFAD microglia ^26,27^, we showed that 5xFAD microglia expressed lower levels of P2RY12 and TMEM119 (Fig. 4B-C). Furthermore, we demonstrated that this decrease in homeostatic markers is dependent on both amyloid pathology and aging. While plaque-associated microglia expressed the lowest levels of both P2RY12 and TMEM119 in all age groups, plaque-distal microglia in 5xFAD mice exhibited a graded decrease in expression of P2RY12 and TMEM119 with aging (Fig. 4B-C). Levels of P2RY12 in 5xFAD plaque-distal microglia were similar to WT microglia at 4 months but decreased to levels comparable to plaque-associated microglia at 9 months. While not as pronounced, the expression of TMEM119 in plaque-distal microglia showed a similar age-related decrease.

**Fig. 4:**
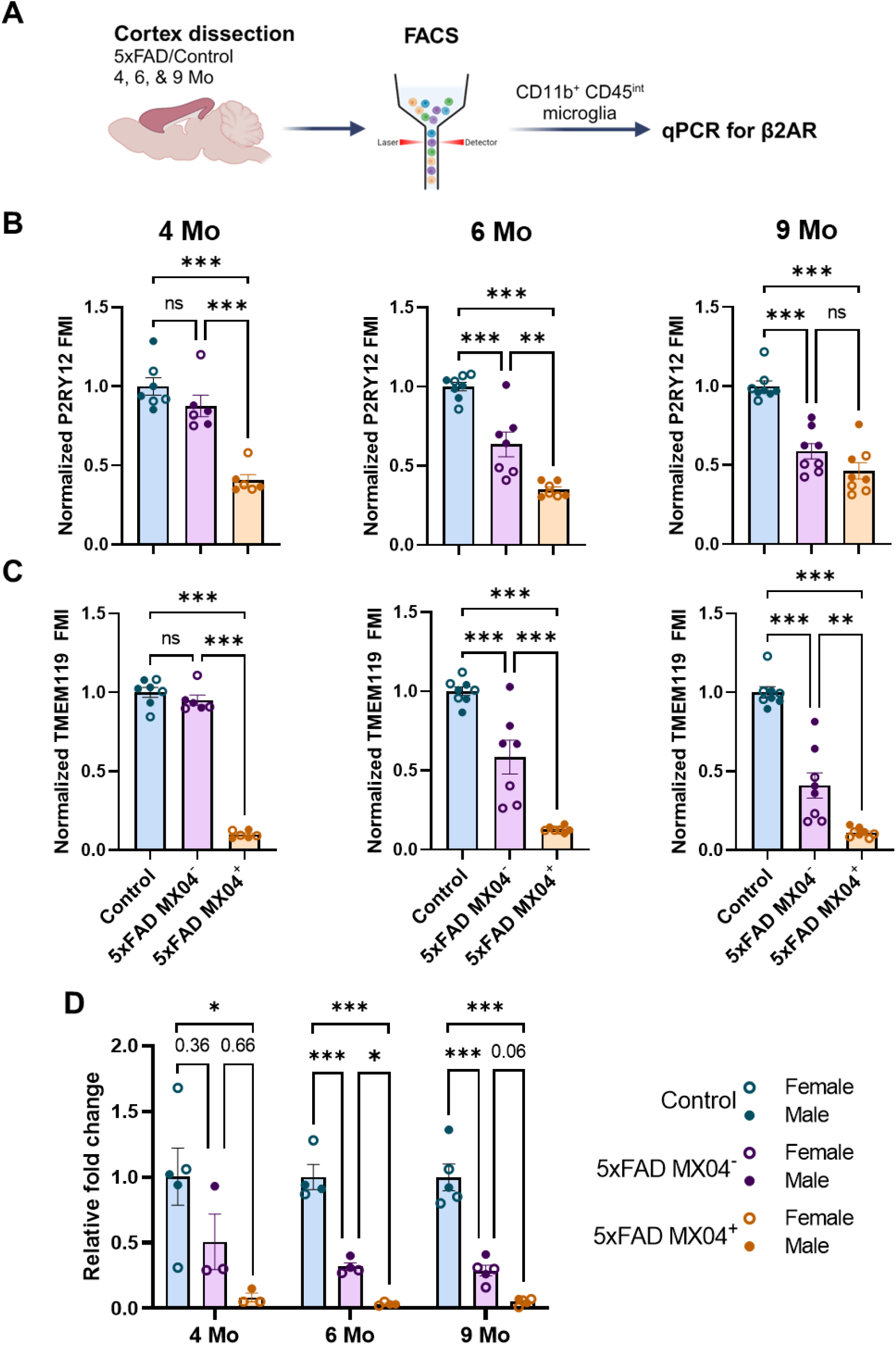
Amyloid pathology and age-dependent synergistic loss in microglial homeostatic signature and β2AR expression in 5xFAD mice. Experimental paradigm for microglia isolation and analysis (**A**). Expression of microglial homeostatic marker P2RY12 (**B**) and TMEM119 (**C)**. Expression of β2AR in isolated microglia measured by qPCR (**D**). *n = 7-9 for FACS, n=3-5 for qPCR, one-way ANOVA with Bonferroni post-hoc correction, *p<0.05, **p<0.01, ***p<0.001*.

We then asked whether downregulation of homeostatic markers correlated with lower microglial β2AR expression. Using qPCR, we detected a more than 10-fold decrease in β2AR expression in plaque-associated MX04^+^ microglia compared to WT microglia at early ages (4 months), and these decreased levels were maintained at 6 and 9 months (Fig. 4D). Plaque-distal MX04^−^ microglia displayed intermediate levels of β2AR expression, which seemed to decrease slightly with age.

### Microglial responsiveness to β2AR stimulation is unaffected by amyloid pathology but decreases with age

Because of the substantial downregulation of β2AR expression in 5xFAD microglia, we examined whether β2AR stimulation could still affect their dynamics *in vivo*. Because *in vivo* imaging necessitated the use of animals haploinsufficient for CX3CR1, we first verified that microglia in this model showed the same downregulation of homeostatic markers and β2AR with age and amyloid pathology. Indeed 5xFAD/ CX3^G/+^ microglia showed a similar pattern of loss of P2Y12 and TMEM119 (Supplementary Fig. 4 vs. Fig. 4). β2AR expression was lost early in plaque-associated microglia but plaque-distal microglia expressed control levels of β2AR at 4 months which decreased to the levels seen in plaque-associated microglia by 6 months (Supplementary Fig. 4C), suggesting that amyloid pathology also affected β2AR expression in CX3CR1 haploinsufficient animals. To test whether these microglia still responded to pharmacological β2AR stimulation, we implemented our previously established experimental paradigm ^23^ in which anesthetized mice were injected with the brain-permeant β2AR selective agonist clenbuterol during imaging sessions to allow for intra-animal comparisons of pre- and post-β2AR stimulation. Animals were pre-dosed with the brain-impermeant β2AR antagonist nadolol at least 1 h before clenbuterol dosing to account for indirect effects of clenbuterol on cardiorespiratory systems during imaging. Clenbuterol treatment caused a rapid retraction of microglial processes (magenta) which was sustained over the next 30 min of imaging (Fig. 5A), resulting in a significant decrease in microglia surveillance of the parenchyma in 4- and 6-month-old mice of both genotypes (Fig. 5B-C). While effects were larger in WT mice, both plaque-distal and plaque-associated microglia also showed significant responses to clenbuterol despite their low β2AR mRNA levels (Supplementary Fig. 4C). Interestingly, clenbuterol did not alter surveillance in either genotype at 9 months (Fig. 5D), despite higher expression of β2AR mRNA in control animals (Supplementary Fig. 4C). Sholl analysis revealed that microglial ramification tracked their surveillance (Fig. 5E-F). This shows that the sensitivity of microglia to pharmacological β2AR stimulation may not scale with mRNA expression of the receptor but is modulated by age. It also suggests that both plaque-distal and plaque-associated microglia may still be modulated my pharmacological intervention in later stages of AD as evidenced by the results in 6-month-old mice in this experiment.

**Fig. 5:**
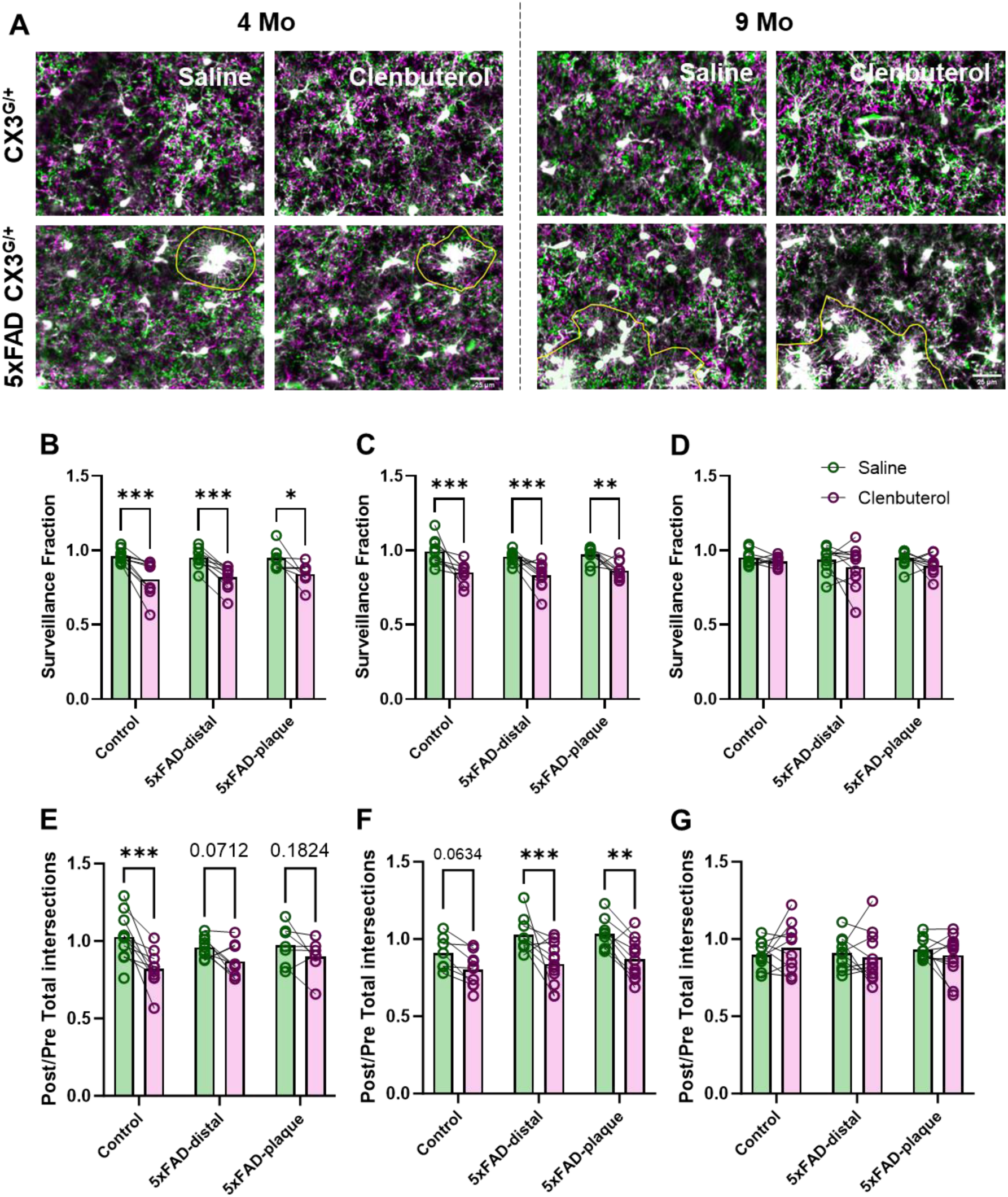
β2AR stimulation with CN decreases microglia surveillance regardless of amyloid pathology but becomes ineffective with age in V1. Representative *in vivo* two-photon time projected images from CX3CR1^GFP/+^(Control) and CX3CR1^GFP/+^5xFAD^+/-^(5xFAD) mice with saline or clenbuterol treatment (**A**). Extended processes in green, retracted processes in magenta. Quantification of microglial surveillance fraction (area of image covered by microglia post/pre) in Saline (green) and Clenbuterol (magenta) treatment groups in Control and 5xFAD mice (**B:** 4 months**; C:** 6 months**; D:** 9 months). Both control and 5xFAD microglia retract their processes in response to clenbuterol at 4 and 6 months but not at 9 months, quantified by taking ratio of microglia total intersections from Sholl analysis post/pre treatment (**E:** 4 months**; F:** 6 months**; G:** 9 months). *n = 9-11, Two-way ANOVA with Bonferroni (B-D) or Tukey (E-G) correction, *p<0.05, **p<0.01, ***p<0.001; Scale bar: 25μm*

### Inhibition of microglial β2AR signaling accelerates amyloid pathology in 5xFAD mice

To determine whether blocking microglial β2AR signaling exacerbates amyloid pathology, we employed both genetic and pharmacological approaches to explore the impact of prolonged absence of microglial β2AR signaling. To selectively ablate β2AR expression in microglia, we treated tamoxifen-inducible microglial-specific β2AR knock-out mice ^31^ on a 5xFAD background (5xFAD CX3CR1-Cre^ERT^ β2AR-flox) with tamoxifen from P41-P45, prior to senile plaque formation. PCR for floxed and excision alleles on isolated microglia from tamoxifen-treated animals confirmed successful gene excision of β2AR (Supplementary Fig. 6). Genetic deletion of microglial β2AR prior to plaque deposition aggravated amyloid pathology and neuritic damage in females with robust increases in LAMP1^+^ area in ACC and V1, as well as trends towards elevation of 6E10^+^ plaque load in both areas and increased Iba1 immunoreactivity in ACC (Fig. 6B-E; 11A-D). Although microglial β2AR ablation in males did not result in significant effects, we observed similar trends towards worsening amyloid pathology in the ACC, where amyloid deposition occurs early (Supplementary Fig. 5I-L). Interestingly, while females showed no change in TH^+^ projections in either area, these were decreased in both areas in males (Supplementary Fig. 5K, O), suggesting a possible sex-specific interaction between β2AR signaling and TH^+^ fiber degeneration.

**Fig. 6:**
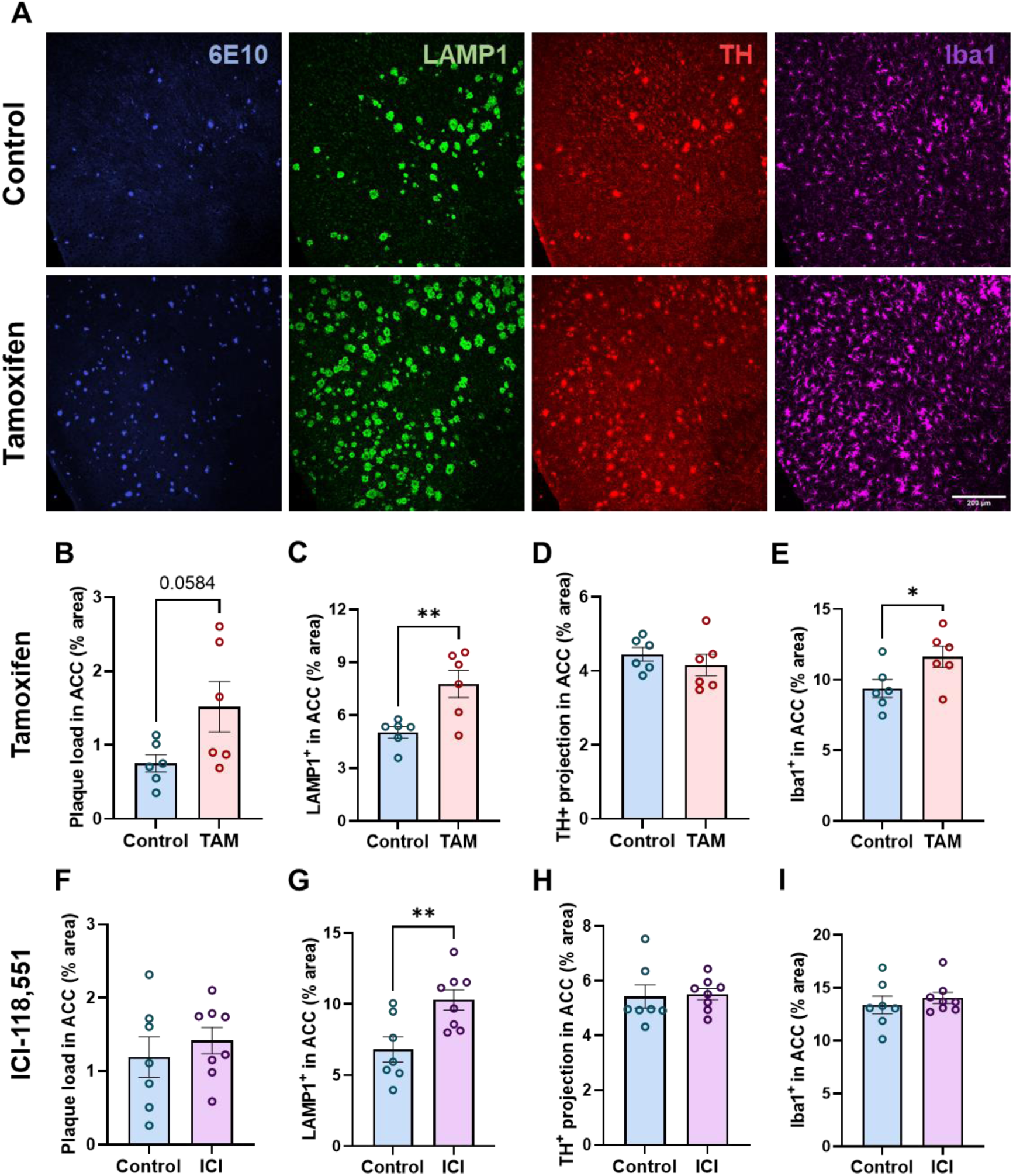
Both genetic deletion of microglial β2AR and prolonged exposure β2AR antagonist accelerates pathology progression in female 5xFAD mice. Representative 20x confocal images of the ACC immunolabeled with 6E10 (plaque), LAMP1 (plaque-associated neuritic damage), TH (putative NE projections) and Iba1 (microglia) (**A**). Quantification of plaque load, neuritic damage, TH^+^ projections and microglia activation in the ACC of control and treated female 5xFAD animals, with microglia-specific deletion of β2AR (**B-E**) or prolonged 1-month treatment with ICI-118,551, a β2AR-specific antagonist (**F-I**). *n = 6-8, Student t-test; *p<0.05, **p<0.01; Scale bar: 200μm*

To complement this genetic approach, we carried out a pharmacological treatment of 1-month ICI-118,551 administration via subcutaneous osmotic pump to selectively block β2AR. Only female 5xFAD mice were included for this experiment because males did not tolerate implanted osmotic pumps for a prolonged period of time. Similar to what we observed with β2AR deletion, chronic inhibition of β2AR dramatically accelerated neuritic damage, without altering senile plaque load, TH^+^ projections or Iba1+ area (Fig. 6F-I).

### Chronic microglial β2AR stimulation attenuates amyloid pathology in 5xFAD mice

After observing the robust effects of single-dose β2AR stimulation on microglia despite the decrease in microglial β2AR expression and a lack of response to anesthesia which is modulated by endogenous NE levels, we next investigated the impact of chronic microglial β2AR activation in 5xFAD mice. After 1 month of daily intraperitoneal clenbuterol injection during their active phase which started at 3 months of age, both male and female 5xFAD mice displayed reduced amyloid plaque load and associated neuritic damage evidenced by lower levels of both 6E10 and LAMP1 immunolabelling (Fig. 7B-C, F-G; Supplementary Fig. 7A-B, E-F), although these did not reach significance except for LAMP1 decreases in males. Quantification of TH^+^ projections revealed no differences between treated and untreated animals (Fig. 7D, H; Supplementary Fig. 7C, G), suggesting that supplementing 5xFAD mice with exogenous β2AR stimulant does not correct the degeneration of LC projections. Iba1+ area was also unchanged with clenbuterol treatment, except for a mild but significant decrease in female ACC (Fig. 7E, I; Supplementary Fig. 7D, H), warranting further investigation on the effects of clenbuterol treatment on microglia characteristics. Previous research showed that NE depletion decreases microglia recruitment to Aβ plaque, which is rescued after only 24 h with NE-precursor L-threo-DOPS treatment ^9^. Thus, we examined whether the decrease in plaque load with chronic activation of microglial β2AR could be explained by enhanced microglia migration towards and phagocytosing Aβ plaque. However, we did not observe any change in microglia recruitment to Aβ plaque in response to chronic β2AR stimulation evidenced by similar microglia coverage per plaque area (Iba1/6E10) between control and treated mice (data not shown).

**Fig. 7:**
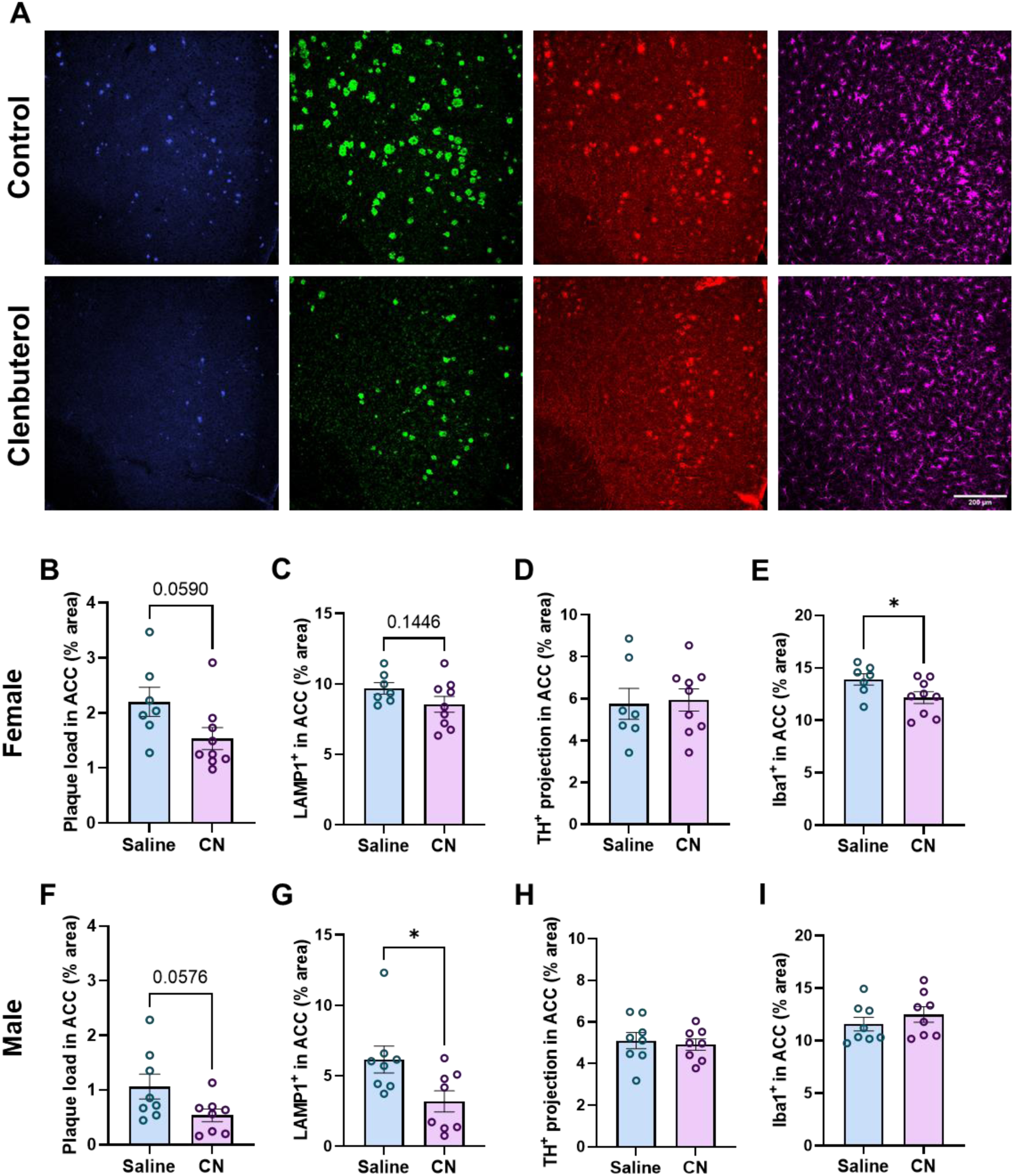
Prolonged exposure to β2AR agonist attenuates amyloid pathology and associated neuritic damage in the ACC in 5xFAD mice. Representative 20x confocal images of the ACC immunolabeled with 6E10 (plaque), LAMP1 (plaque-associated neuritic damage), TH (putative NE projections) and Iba1 (microglia) (**A**). Quantification of plaque load, neuritic damage, TH^+^ projection and microglia activation in cortex of control and treated 5xFAD females (**B-E**) and males (**F-I**) with prolonged 1-month treatment with Clenbuterol, a potent β2AR agonist. *n = 8-9, Student t-test; *p<0.05, **p<0.01. Scale bar: 200μm*

## Discussion

In the present study, we report a systemic characterization of endogenous NE-releasing LC neurons, their cortical projections, levels of released NE as well as microglial sensitivity to NE in the brains of 5xFAD mice of 4, 6, and 9 months of age, representing early, mid and late amyloid pathogenesis respectively. We found early loss of NE cortical fibers which correlated with reduced cortical NE levels and at later ages reduced cortical NMN levels. LC neuronal loss was only observed in advanced stages of the disease. Isolating microglia from 5xFAD and control mice revealed a striking loss of microglial mRNA expression of β2AR, the receptor chiefly responsible for translating direct NE signaling to microglia ^23,24^. This loss was especially profound in microglia directly associated with plaques even at young ages. The synergistic degeneration of the LC-NE system and microglial responsiveness to NE resulted in impaired microglial dynamics in 5xFAD mice. Interestingly, aging alone, in the absence of amyloid pathology, also diminished microglial response to endogenous NE. Most importantly, we demonstrated the potential of microglial β2AR as a specific drug target, showing that pharmacological targeting of β2ARs could alter microglial behavior and attenuate plaque deposition despite the decreases in expression of the receptor on microglia. On the other hand, inhibition of this receptor, or specific deletion of the receptor in microglia, worsened plaque pathology, implicating loss of direct signaling of NE to microglia in detrimental outcomes in AD. Our findings suggest the potential to leverage microglial β2AR signaling for AD disease-modifying therapies (working model Supplementary Fig. 8).

### Degeneration of the endogenous noradrenergic system 5xFAD mice

Our characterization of changes in the endogenous NE system shows that 5xFAD mice show changes in the noradrenergic system at both early and late stages of amyloid pathology. Characterization of TH^+^ LC neurons revealed a significant reduction in cell number with concomitant increases in soma size only in 9-month-old 5xFAD mice compared to WT littermates (Fig. 1). In agreement with our results, Kalinin et al., reported no significant differences in TH^+^ LC neuron number between 6-monthold 5xFAD and WT mice ^14^. Other studies using different experimental designs in models where amyloidosis develops more slowly have collectively shown significant reduction in LC neuron number from 9-12 months of age ^28,42–45^. Associated hypertrophy of LC neurons was also noted in both APP/PS1 ^43^ and 5xFAD ^14^ mice. While our observed increase in LC neuron size occurred at a later age than reported by Kalinin et al., ^14^, it seems that amyloid pathology can impair the structural integrity of LC neurons as well as lead to their degeneration. However, the presence of dense core plaques themselves is unlikely to be the driving factor of LC neurodegeneration as the LC is devoid of these senile plaques (Supplementary Fig. 1). It is possible that the combination of neuroinflammation evidenced by increased Iba1 and GFAP expression (Supplementary Fig. 1) and vulnerability of LC neurons ^46^, either inherently or through the earlier loss of their projections, are chiefly responsible for LC neuronal loss and hypertrophy.

TH^+^ nerve fibers appear to degenerate early in the brains of 5xFAD mice (Fig. 2), and may be the substrate for altered endogenous NE signaling in AD at a time when LC neuronal cell bodies are still unaffected. The decrease in TH^+^ projections was observed in ACC at 4 months and in V1 at 6 months of age, coinciding with the start of substantial Aβ plaque deposition and increases in neuroinflammation in each respective region (Fig. 2). Similarly, Cao et al., reported significant reduction in TH^+^ area across different cortical and subcortical structures in APP/PS1 ^28^. However, in 5xFAD mice we did not observe further loss of cortical TH^+^ nerve fibers as pathology progressed, suggesting that compensatory mechanisms may prevent further degeneration as plaque load and inflammation increase in the disease. Interestingly, at 9 months of age, WT mice show similar losses in TH^+^ fibers, suggesting that aging may affect fiber degeneration as potently as amyloid pathology (Fig. 2). Though Cao et al., also reported gradual loss of TH^+^ projections in WT mice with aging (3-versus 12- and 18-month-old); at 12 months of age, APP/PS1 mice still exhibited more pronounced degeneration of TH^+^ fibers ^28^. This discrepancy is likely due to differences between the models as 5xFAD mice develop amyloid pathology early with senile plaque presence at ∼2-month and plateaus at ∼9 months ^29,39^, while APP/PS1 mice exhibit sparse plaque deposition at ∼5-6 months and slow disease progression with plaque load peaking at ∼20 months of age ^47^.

Since TH^+^ projections represent axons of both noradrenergic and dopaminergic neurons, we performed ELISA on flash frozen cortices to examine levels of total NE produced and its metabolite NMN, which is the product of synaptic COMT-mediated degradation of NE ^40,41^. Interestingly, we observed an early reduction in total NE, but not NMN in 4-month-old 5xFAD compared to WT mice (Supplementary Fig. 2). With aging, levels of total NE were comparable between 5xFAD and WT mice, similar to a previous report in aged APP/PS1 and WT mice ^28^; whereas NMN levels in 5xFAD mice were lower than that in WT mice although the effect was not significant (Supplementary Fig. 2). We also observed an overall trend of age-dependent increase in cortical NA content, peaking at 6-month-old, although no statistical comparisons were made because animals of different age groups were harvested on different days. Similar findings of age-related NE elevation have been reported in both mice ^48^ and rats ^49–51^. Taken together, our findings suggest two important notions that warrant further investigation. First, it appears that there might be many different compensatory mechanisms that try to normalize NE cortical levels at different stages of amyloidosis. For instance, early loss of NE projections in the frontal cortex may lower NE release but compensatory electrophysiological changes at the synapse to facilitate NE release or increase NE sensitivity may normalize these levels later in the disease. Secondly, even though both aging and advanced amyloid pathology lead to comparable degeneration of NE nerve fibers (Fig. 2), amyloid pathology might impair NE release (Supplementary Fig. 2), dampening the impact of NE in aged AD brains. Previous research suggests that increased activity of remaining LC neurons in AD brains ^52,53^ and increased excitability of LC axons in the frontal cortex during aging ^50,51^ might compensate for the reduction in cortical NE innervation. BDNF is shown to be crucial for maintenance of cortical NE axonal branching and plasticity during aging ^51^. However, both protein and mRNA levels of BDNF in the brain decrease in AD compared to healthy aging ^54–58^, suggesting that BDNF might be an important regulator of synaptic plasticity to influence NE release, which is perturbed in advanced AD pathology.

### Changes in microglial NE sensitivity in 5xFAD mice

In parallel with changes in adrenergic neurons, microglia also showed changes in their sensitivity to NE. Microglia isolated from 5xFAD brains showed an amyloid pathology-dependent downregulation of β2AR mRNA levels, in line with single-cell transcriptomic data that shows lower β2AR expression in the DAM cluster ^26^. Interestingly, plaque-associated microglia showed very low levels of β2AR mRNA at all ages, suggesting that β2AR is downregulated early in the transition to a DAM phenotype. The fact that plaque-distal microglia showed intermediate levels of β2AR expression suggests that amyloid pathology impacts microglial adrenergic function even in the absence of direct interaction of microglia and amyloid plaques. It is worth noting that for all *in vivo* imaging experiments, we used mice with only one functional copy of CX3CR1, which can alter microglial transcriptomic profiles ^59^ and exert a complex role on amyloid pathology progression, slowing down disease onset but facilitating progression ^60,61^. We thus validated our findings in 5xFAD CX3^G/+^ mice, showing similar downregulation of β2AR in plaque-associated microglia regardless of age and an age-dependent reduction in β2AR levels in plaque-distal microglia, which suggest a more pronounced loss of β2AR distal from plaques in later stages of the disease (Fig. 4 vs. Supplementary Fig. 4)

While microglia in young WT mice responded to lower NE levels during anesthesia with increased surveillance as previously reported ^23,24^ (Fig. 3), we show that microglia in both aged 5xFAD and WT mice no longer respond to anesthesia likely due to the age-related degeneration of NE projection fibers. This is an important result suggesting that adrenergic dysregulation of microglial signaling could contribute to the aging-related dysfunctions of microglia reviewed in ^62^. If NE production does indeed rise with age (as discussed above), these results suggest this increase is not sufficient to compensate for the loss in NE axonal innervations, potentially due to impaired local NE dissemination in the synaptic and extra-synaptic areas. It will be important to monitor patterns of spatiotemporal NE release *in vivo*, possibly using multi-photon imaging with robust new NE sensors ^63^ that can provide more reliable measurements than bulk ELISA-based methods. In line with our observations that both NE projections and microglial β2AR expression decrease with amyloid pathology, 5xFAD microglia, especially those associated with plaques, did not respond to anesthesia starting early in the disease (Fig. 3). Surprisingly, despite their striking downregulation of β2AR mRNA expression, plaque-associated microglia were able to respond to direct stimulation of the β2AR by the agonist clenbuterol (Fig. 5), suggesting that the lack of response to anesthesia was not due to a complete loss of this receptor but rather a combination of lower NE release and lower sensitivity to NE of microglia. This also suggests that residual β2AR function in microglia could be targeted therapeutically with pharmacological interventions even in DAM, presenting a possibility for adrenergic therapy later in the disease. Additionally, aged microglia in both 5xFAD and WT mice no longer responded to B2AR stimulation despite the presence of high levels of β2AR mRNA in aged WT microglia (Fig. 4-5; Supplementary Fig. 4). This shows a lack of correspondence in β2AR mRNA levels and microglial sensitivity likely due to β2AR protein trafficking or receptor binding affinity/kinetics. Unfortunately, proteomic assays for the β2AR are unreliable, making it hard to address changes to this receptor at the protein level. It is important to also note that 9 months of age is not generally considered old for WT animals, and the age-related microglia cluster has been shown to downregulate β2AR expression much later at 18 months ^25^. Thus, it is important to profile both β2AR mRNA and protein expression and address the timescale of changes in their expression patterns.

### Microglial β2AR signaling attenuates amyloid pathology

The role of NE as a potent anti-inflammatory agent has been extensively studied in various rodent models of AD where ablation of LC neurons pharmacologically ^9,10,64^, eliminating their NE synthesis and release ^65,66^, and blocking NE actions with β blockers ^11,12^ all exacerbate plaque load and increase levels of inflammatory cytokines. Here, we propose a mechanism through which NE modulates AD pathology. Specific genetic deletion of microglial β2AR exacerbated amyloid pathology, especially plaque-associated neuritic damage, although the effect only reached statistical significance in females (Fig. 6; Supplementary Fig. 5). Pharmacological β2AR inhibition showed similar results in female mice (Fig. 6; Supplementary Fig. 5). Perhaps surprisingly, given the low mRNA expression of β2AR in 5xFAD microglia, we demonstrated that chronic treatment with β2AR agonist was protective, resulting in trends towards less plaque load and neuritic damage in both sexes (Fig. 7; Supplementary Fig. 7). This may be due to the different regulation of β2AR protein as compared to mRNA in microglia as described above, but this also raises the possibility that β2AR stimulation later in the disease could provide therapeutic benefit. It will be important to define the durations and timing of β2AR stimulation that confer benefits in the context of AD pathology, although the larger effects on neuritic damage in male vs. female mice suggests that early interventions may provide the largest improvements.

Our results open new avenues for future studies to investigate microglial β2AR downstream signaling pathways in disease contexts. β2AR and P2RY_12_ signaling have been proposed to act in a push-pull system and their balance is crucial for a myriad functions of microglia, evidenced in recent research showing their opposing effects on microglial morphology, motility and chemotaxis ^23,24,67,68^. However, both β2AR and P2RY_12_ expression significantly decrease with amyloid pathology, making it difficult to detangle their respective roles in mediating downstream intracellular G-protein signaling pathways. Microglia make contacts with both inhibitory and excitatory synapses to control their excitability and plasticity ^68–70^, thus, it is important to characterize microglia-neuron interaction that might be altered in AD in response to diminishing NE or purinergic signaling.

It should also be noted that these findings do not exclude the possibility that the NE-β2AR signaling pathway can also indirectly influence microglia functions to modulate AD pathology. Although expressed at much lower levels compared to microglia ^22^, astrocytic β2ARs are necessary for hippocampal long-term memory formation and consolidation ^71–73^, the deterioration of which is a prominent clinical manifestation of AD. Astrocyte and microglia activity are intimately linked in regulating brain development, synaptic transmission, and glial functional states ^74^, thus, it is possible that β2AR signaling triggers a positive-feedback loop shifting astrocyte and microglia to a reactive state. Nonetheless, indirect signaling could positively contribute to the potential therapeutic benefits of chronic β2AR stimulation.

A vast literature describes sex-specific signaling in AD and its mouse models, making it possible that microglial signaling through the β2AR may lead to different outcomes in males and females. However, we believe that the smaller not-quite-significant effects observed in male mice after genetic β2AR deletion from microglia may be due to the inherent sparse Aβ plaque deposits at this age in males ^29,39^ in combination with the use of CX3CR1 haploinsufficient animals, in which the onset of senile plaque formation is delayed ^60,61^, making it harder to quantify changes in pathology. This would implicate disease stage, rather than sex in the different results obtained in the two sexes. Unfortunately, we could not replicate these results pharmacologically, because male mice could not tolerate long-term osmotic pump implantation. This sex-specific vulnerability to surgery may be interesting and should be explored in the future.

Pharmacological stimulation of β2AR receptors through daily injections caused similar changes in male and female mice, suggesting that β2AR signaling can be harnessed for therapy in both sexes. However, sex-specific differences in microglial β2AR signaling in AD should continue to be explored.

## Methods

### Experimental animals

All animal procedures were reviewed and approved by the University Committee on Animal Resources of the University of Rochester Medical Center and performed according to the Institutional Animal Care and Use Committee and guidelines from the National Institute of Health (NIH). Animals were housed in a 12-hour light/12-hour dark cycle with ad libitum access to standard rodent chow and water. Mice used in long-term clenbuterol treatment experiment were housed in a reverse light/dark cycle, and treatment was given 4 h into the dark cycle. Male and female B6.Cg-Tg(APPSwFlLon,PSEN1*M146L*L286V)6799Vas/Mmjax mice (5xFAD, ^29^) were obtained from JAX (stock no. 034848) and maintained at the University of Rochester vivarium. For two-photon microscopy, 5xFAD mice were crossed with homozygous CX3CR1-GFP reporter mice (JAX stock no. 005582 ^30^, to generate CX3CR1^GFP/+^ 5xFAD and CX3CR1^GFP/+^ littermate controls. For selective genetic deletion of microglial β2AR; 5xFAD, CX3CR1Cre^ERT^ (JAX stock no. 021160) and β2AR-flox ^31^ (Karsenty laboratory, courtesy of the Rosen laboratory) were crossed to generate mice that are heterogenous for 5xFAD CX3CR1Cre^ERT^ and homozygous for β2AR-flox. All mice were derived from and maintained on a C57/Bl6 background.

### Pharmacological agents

Fentanyl cocktail comprised fentanyl (0.05 mg/kg), midazolam (5.0 mg/kg) and dexmedetomidine (0.5 mg/kg) premixed in saline and was injected intraperitoneally (i.p.) for anesthetized two-photon imaging sessions.

Nadolol (10 mg/kg i.p.; Sigma, 42200-33-9) was administered for two-photon imaging. Clenbuterol (1 mg/kg i.p.; Sigma, 21898-19-1) was administered for two-photon imaging, while 2 mg/kg i.p. daily (5 days/week) for 1 month was administered to assess the long-term impact of β2AR agonist treatment. Both nadolol and clenbuterol were dissolved in saline. ICI-118,551 (Sigma, 72795-01-8) were dissolved in DMSO (Sigma, 67-68-5) and administered by mini-osmotic pump (Alzet, model 2002) for 1 month at a dosage of 10 mg/kg/day. Pumps were replaced once after the first 15 days of the treatment period.

Tamoxifen (Sigma, 10540-29-1) was dissolved in corn oil (20 mg/mL) and administered i.p. for 5 days (75 mg/kg) in experiments using 5xFAD CX3CR1-Cre^ERT^ β2AR-flox mice.

### Brain harvesting

In some cases, mice were injected 24 hours before sacrifice with Methoxy-X04 (MX04, i.p., 4mg/kg, Tocris Biosciences), a brain-permeable Aβ fluorescent marker ^32,33^. On the day of brain harvesting, animals were deeply sedated with sodium pentobarbital overdose (Euthasol 1:10; Virbac) and perfused intracardially with 0.1M phosphate buffer saline (PBS). After perfusion, hemispheres were separated: one was immediately submerged in fixative solution (4% paraformaldehyde (PFA), pH 7.2 in PB, 4°C) to be used for immunofluorescence experiments, and cortex was dissected from the other hemisphere and separated into one-half for Fluorescence activated cell sorting (FACS) and one-half flash-frozen in cold isopentane for ELISA.

### Microglia isolation and RNA analysis

#### Fluorescence activated cell sorting (FACS)

The cortical tissue was homogenized in 3 mL FACS buffer (1X PBS + 0.5% BSA). Homogenates were filtered through a 70 µm cell strainer into a 15 ml tube containing 3 ml FACS buffer. The strainer was washed with an additional 3 ml of FACS buffer, and the cell suspensions were centrifuged at 400 x g for 5 min at 4°C. The supernatants were discarded, and the remaining pellets were resuspended in 40% Percoll (Cytiva) (diluted in PBS), then centrifuged at 400 x g for 30 min with no braking. After removing the supernatants, the pellets were resuspended in 90 µL FACS buffer with Fc block (4G2, 1:90, BioLegend) and transferred to a 96 well-plate. After a 15 min incubation with Fc block at 4°C, the following antibodies were added in a 10 µl master mix: CD11b-FITC (M1/70, 1:400, 1:400, Biolegend), CD45-APC/Cy7 (30F11, 1:400, Biolegend), P2Ry12-APC (S16007D, 1:50, Biolegend) & TMEM119-PE (106-6, 1:500, Abcam). The latter two cell surface molecules are considered homeostatic microglial markers ^26,27^. The plate was then incubated for 30 mins at 4°C in the dark. The samples were washed once with FACS buffer and transferred to 5 ml tubes containing 7AAD (Invitrogen) such that its final dilution was 1:80. Appropriate fluorescent-minus-one (FMO) and single-stained bead controls (Ultracomp eBeads, Invitrogen) were prepared in tandem with samples. After excluding debris, doublets, and dead cells, CD11b+/CD45int was used to gate for microglia on a FACSAria II (BD). MX04+ and MX04-microglia were sorted. All events were recorded, and data were analyzed with FCS Express 7 (DeNovo Software).

#### RNA isolation and quantitative PCR

Sorted cells were collected in 300 μL RLT Buffer (Qiagen) and total RNA was isolated using the RNeasy Plus Micro Kit (Qiagen). RNA concentration was determined with the Nanodrop ND-1000 spectrophotometer (NanoDrop) and RNA quality assessed with the Agilent Bioanalyzer (Agilent). Samples with at least 0.5 ng RNA were amplified with the NuGEN Ovation RNA Amplification Kit (Tecan) per manufacturer’s recommendations. The quantity and quality of the subsequent cDNA was determined using the Qubit Fluorometer 3.0 (Invitrogen) and the Agilent Bioanalyzer. Quantitative PCR was run in a 96-well plate format on a QuantStudio Q3 system, with 2 technical replicates per sample. For each well, a final volume of 10 μl reaction containing 50 ng of cDNA sample, TaqMan Fast Advanced Master Mix (Applied Biosystems) and Taqman Gene Expression Assays for GAPDH and β2AR was loaded. Samples were denatured at 95°C for 5 min, followed by 40 cycles of denaturing at 95°C for 30 s, annealing at 60°C for 20 s and extension at 72°C for 30 s. Fold changes were determined with the 2^-ΔΔCt^ method.

### Confirmation of β2AR excision on sorted microglia

Sorted cells were collected in tubes containing 300 μL of 50 mM NaOH, boiled for 10 min at 100°C, vortexed and then 75 μL 100mM Tris pH 6.8 was added to isolate DNA. A total of three tamoxifen-treated and three untreated animals were included in these confirmation experiments. Isolated DNA was run through two PCR protocols: first, the PCR for confirmation of floxed allele presence (550 bp product: Forward (Fw), CCAAAGTTGTTGCACGTCAC; Reverse (Rv), GCACACGCCAAGGAGATTAT); and second, excision confirmation (∼800 bp product: Fw, CCAAAGTTGTTGCACGTCAC; Rv, AAGAAAGAGGAGGGGCTGAG). Similar to our previous report ^23^, we confirmed that the floxed allele PCR product was no longer present in tamoxifen-treated mice; however, there was a certain degree of leakiness in our Cre expression evidenced by the presence of excision product in both treated and untreated mice (Supplementary Fig. 6)

### ELISA

Cortical tissue from half of one hemisphere was weighed and homogenized at 10% w/v in 0.01N HCl. To avoid degradation of NE, tissue was kept on ice and covered in aluminum foil throughout the procedure whenever possible. NE and its metabolite normetanephrine (NMN) levels were measured utilizing respective ELISA kits (Rocky Mountain Diagnostics) per manufacturer’s recommendations, with 2 technical replicates per sample. Plates were read with a Microplate Absorbance Reader (Bio-Rad). It should be noted that data from tissue collected for different age groups were analyzed separately as cortices were collected at different times leading to variability in measurements between age groups.

### Immunofluorescence

Half-brains were fixed overnight in 4% PFA at 4°C, dehydrated in 30% sucrose overnight, and sectioned on a freezing stage microtome into 30 µm thick coronal slices stored in cryoprotectant solution. For immunofluorescence, sections were washed extensively in PBS and blocked with 10% bovine serum albumin (BSA, Sigma, A2153) for 1 h at room temperature (RT).

Sections were immunolabeled for amyloid-beta (Aβ), microglia (Iba1), astrocytes (GFAP), tyrosine hydroxylase (TH), and a widely used marker for neuritic damage LAMP1 ^15,34–36^; in different combinations specified in appropriate figure legends. The following primary antibodies were used: biotin anti-Aβ (clone 6E10, 1:3000, BioLegend), rabbit anti-Iba1 (1:2000, Wako), guinea pig anti-GFAP (1:3,000, Synaptic Systems), mouse anti-TH (1:500, Millipore Sigma) and rat anti-LAMP1 (1:2000, Abcam). Sections were incubated in primary antibodies for 48 h at 4°C. The sections were washed with PBS and incubated in fluorescently labeled secondary antibodies/reagents (Alexa Fluor 405, Alexa Fluor 488, Alexa Fluor 594, streptavidin conjugate 594 and Alexa Fluor 647, Invitrogen; all at 1:1000) for 4 hr at RT, then mounted and coverslipped (Prolong Diamond, ThermoFisher Scientific).

### Confocal microscopy image acquisition and analysis

For assessment of LC neurons, all sections containing LC were selected. For all other experiments, 3 coronal tissue sections that included the anterior cingulate cortex (ACC) and 3 that included the primary visual cortex (V1) were selected. Z-stacks of the region of interest were captured with a Nikon A1R HD confocal microscope using a 20x (Plan Apo VC, 0.75 NA) objective lens. Imaging parameters were kept constant across all sections for each set of immunofluorescent labels. All image analysis was performed using Cellpose 0 with custom trained model and ImageJ FIJI (NIH) with semi-automated custom macros. Experimenters were blind to treatment.

Analysis of LC neuron number and size was done by optimizing the generalist algorithm for cell and nucleus segmentation Cellpose 0. A heterogenous set of 40 LC images was used in the training with the following parameters: cyto2 pretrained model, 250 epochs, 0.1 learning rate and 0.0001 weight decay. The custom model was then applied to all images containing LC to LC neuron number and size, batch processing was done in Napari.

Analyses of amyloid pathology, microglia and astrocyte reactivity, TH^+^ projections and LAMP1 were performed in ImageJ with custom macros. All images were subjected to preprocessing steps including despeckle and background correction. Regions of interest (ROIs) outlining the LC, ACC, and V1 were drawn on maximum-intensity projection of the acquired images. Images were subsequently thresholded and binarized using automated ImageJ thresholding algorithms, which were kept consistent for all images in each experiment. For all markers, the area fraction was calculated as the ratio between the number of pixels above the threshold over all pixels in the ROIs. Due to the artifacts of TH immunoreactivity at Aβ plaques as previously reported ^28,37^, an extra step of subtracting plaque area from the TH channel was performed.

### Cranial window surgery

Animals were anesthetized using the fentanyl cocktail (i.p.) during the cranial window implantation surgical procedure. Body temperature was maintained at 37°C with a heating pad and the animal’s eyes were protected with lubricant ointment. All conducted surgical procedures adhered to aseptic technique. Mice were fixed in a stereotaxic frame; hair was removed and the skull was exposed through a scalp incision. A 3-mm biopsy punch (Integra) was then used to create a circular score on the skull over V1. A 0.5-mm drill bit (FST) was used to then drill through the skull for the craniotomy, tracing the 3-mm score. A 5-mm coverslip attached to a 3-mm coverslip (Warner Instruments) by UV glue (Norland Optical Adhesive, Norland) was then slowly lowered into the craniotomy (3-mm side down). The coverslip was carefully secured with C&B Metabond dental cement (Parkell). A custom headplate produced by emachine shop (http://www.emachineshop.com) (designs courtesy of the Mriganka Sur laboratory, Massachusetts Institute of Technology) was then secured onto the skull with the same dental cement, the rest of which was used to cover any exposed skull and seal the incision site. Mice were administered slow-release buprenex (5 mg/kg subcutaneously for 72 h) and carprofen (5 mg/kg, i.p. every 24 h) and monitored for 72 h postoperatively.

### Two-photon microscopy image acquisition and analysis

A custom two-photon laser-scanning microscope was used for *in vivo* imaging (Ti:Sapphire, Mai-Tai, Spectra Physics; modified Fluoview confocal scan head, 20x water immersion objective lens, 0.95 NA, Olympus). Excitation for fluorescence imaging was achieved with 100-fs laser pulses (80 MHz) at 920 nm for GFP and 770 nm for MX04 with a power of ∼40-50 mW measured at the sample. Fluorescence was detected using a photomultiplier tube with a 580/180 bandpass filter (GFP, microglia) and 460/80 filter (MX04-labeled plaque). Mice were injected with MX04 (i.p., 4 mg/kg) 24 hr prior to the first imaging session and given additional doses once every 5 days (i.p., 1 mg/kg).

For anesthetized imaging sessions, mice were anesthetized with the fentanyl cocktail. During and post-imaging, body temperature was maintained at 37°C with a heating pad and the animal’s eyes were protected with lubricant ointment. Time-lapse imaging was carried out at 5 min intervals over 1.5 h, 45-60 μm z-stack depth at 1 μm step size at each time point, 4x digital zoom. For β2AR stimulation, clenbuterol or saline was administered 30 min into the imaging session, allowing for intra-animal comparisons of pre-(first 30 min) and post-(last 30 min) stimulation. To limit peripheral effects of clenbuterol, nadolol was given 1 h before the imaging session. Prior to awake imaging sessions, mice were allowed to habituate on the running wheel over at least three sessions. Mice were head-fixed on the apparatus for 30 min in the first session and for increasing amounts of time in the subsequent sessions to a maximum of 1.5 h. Time-lapse recordings were collected as described above for 1 h. For repeated imaging, blood vessels were used as gross landmarks and stable microglia were also used as fine landmarks to re-identify the correct region for imaging. Image analysis was done offline using ImageJ, Ilastik and MATLAB with custom algorithms.

All images are subjected to preprocessing steps in ImageJ as previously described ^23,38^ and macros are available at https://github.com/majewska-lab. Due to high motion artifact in images acquired from awake imaging sessions, principal component analysis (PCA) was performed, and images were reconstructed using a quantitatively determined number of components with a custom MATLAB script to remove high frequency image noise.

For assessment of microglia dynamics, microglia surveillance was quantified, representing how much of the parenchyma microglia survey over time. For automated detection of microglial processes, the image classification and segmentation software, Ilastik, was used. To train for pixel classification, microglia processes and somas were manually traced. Appropriate thresholding and size exclusion criteria were applied for object classification. Outputs of microglial processes were binarized in ImageJ for microglia surveillance analysis. The surveillance index was calculated by dividing the number of binarized microglia pixels by all pixels in the maximum projection of all timepoints. For β2AR stimulation experiment, the surveillance fraction was calculated as the ratio between post and pre-treatment surveillance index.

For assessment of microglia morphology, Sholl analysis was performed on at least 2 microglia per animal in an imaging session to quantify degree of ramification. All microglia processes were manually traced on 2D z-maximum intensity projections, binarized and analyzed with an automated ImageJ Sholl Analysis plug-in. The maximum and total number of intersections were used for statistical analyses.

### Statistical analysis

Data organization and summary were carried out in RStudio v4. All statistical analyses and graphing were performed in Graphpad Prism v9. Comparisons between two genotypes/treatments were made using the Student’s *t*-test. Comparisons among more than two groups were done using either one-way or repeated-measure ANOVA when suitable with appropriate post-hoc correction. Detailed statistics are provided in the appropriate figure legends. All data points represent individual animal averages and are presented as mean ± SEM.

## Acknowledgements

We thank Jean M. Bidlack for her advice on the design of experiments with pharmacologic agents. We thank Berke Karaahmet, Elizabeth Plunk, Mark Stoessel, Lee Trojanczyk, and Nisha Arya for technical assistance with tissue procurement and processing. We thank the staff of Center for Advanced Light Microscopy and Nanoscopy (CALMN; University of Rochester Medical Center) for image acquisition discussions. We also thank the Flow Cytometry Core (FCC; University of Rochester Medical Center) for technical assistance. We thank the Genomics Research Center (GRC; University of Rochester Medical Center) for processing samples for RNA extraction and quality assessment. This research was supported by the Alzheimer’s Association AARG-NTF-19-619116 (AKM), University of Rochester Medical Center’s Del Monte Institute for Neuroscience Pilot Program (AKM, KO), a University of Rochester Goodman award (LL), and an AD supplement to NIH R01 NS114480 (AKM, KO).

## Author contributions

LL conceived the study in consultation with AKM and MKO. LL designed, conducted, and carried out data analyses for most of the experiments in this study with the following contributions from co-authors. AF performed Sholl analyses for all *in vivo* imaging data and assisted with FACS experiments. CL, HL and KKP assisted with histology experiments pertinent to β2AR agonist and antagonist treatments. LL and AKM wrote the first draft of the manuscript. All authors reviewed and contributed to the final version.

## Competing interests

The authors declare no competing interests.

## Data availability

The raw data supporting the findings of this study and training models for image analysis will be made available from the corresponding author upon reasonable request from any qualified researcher.

---

**Supplementary Fig. 1:**
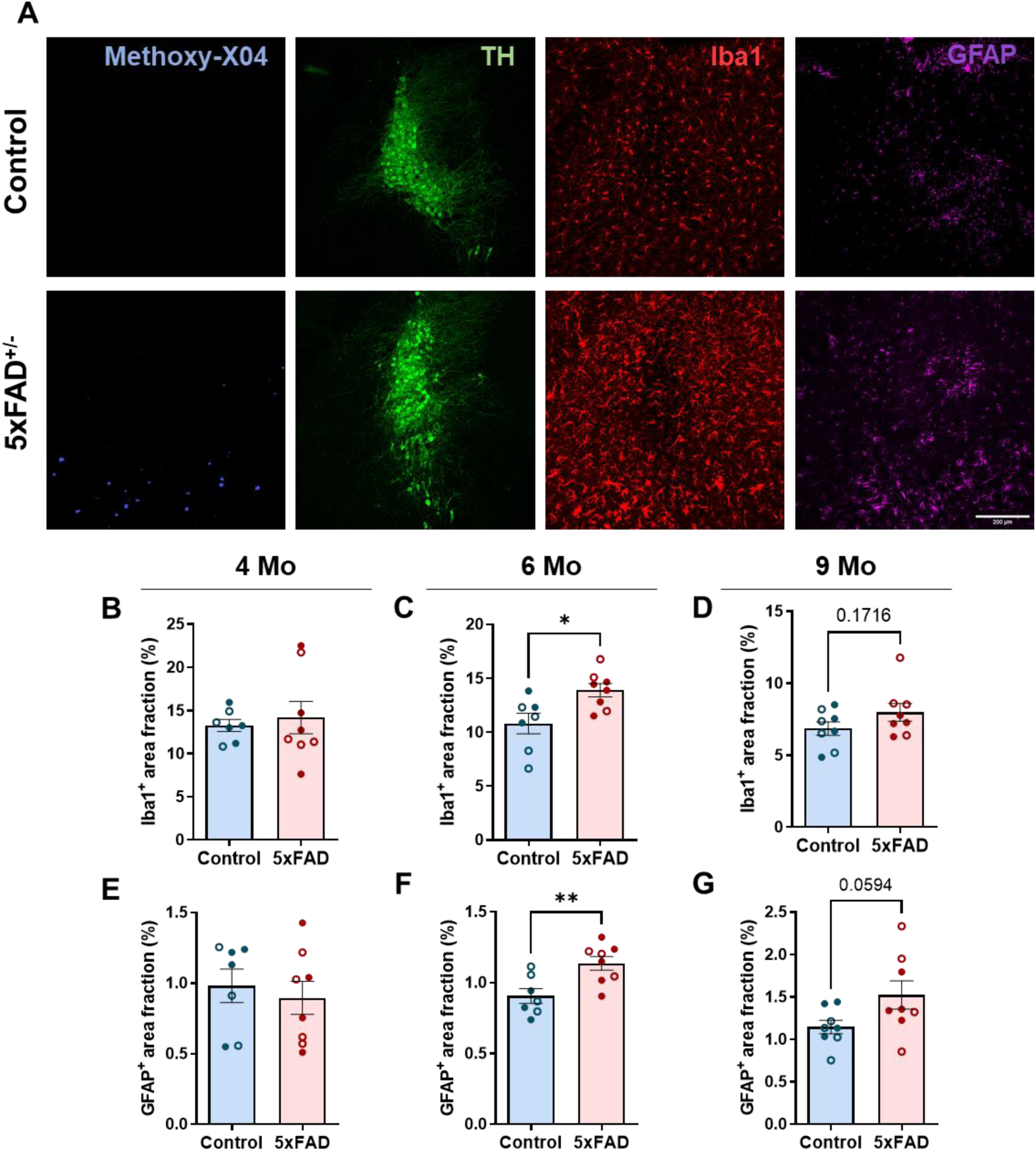
exhibit glia activation during advanced disease stages in 5xFAD mice. Representative 20x confocal images showing neuritic plaques (Methoxy-X04, blue), LC neurons (TH, green), microglia (Iba1, red), and astrocytes (GFAP, magenta) in the LC (**A**). Quantification of Iba1^+^(**B-D**) and GFAP^+^ (**E-G**) area fraction as proxy for glia activation across 3 age groups. *n = 7-9, Student t-test; *p<0.05, **p<0.01. Scale bar: 200μm*.

**Supplementary Fig. 2:**
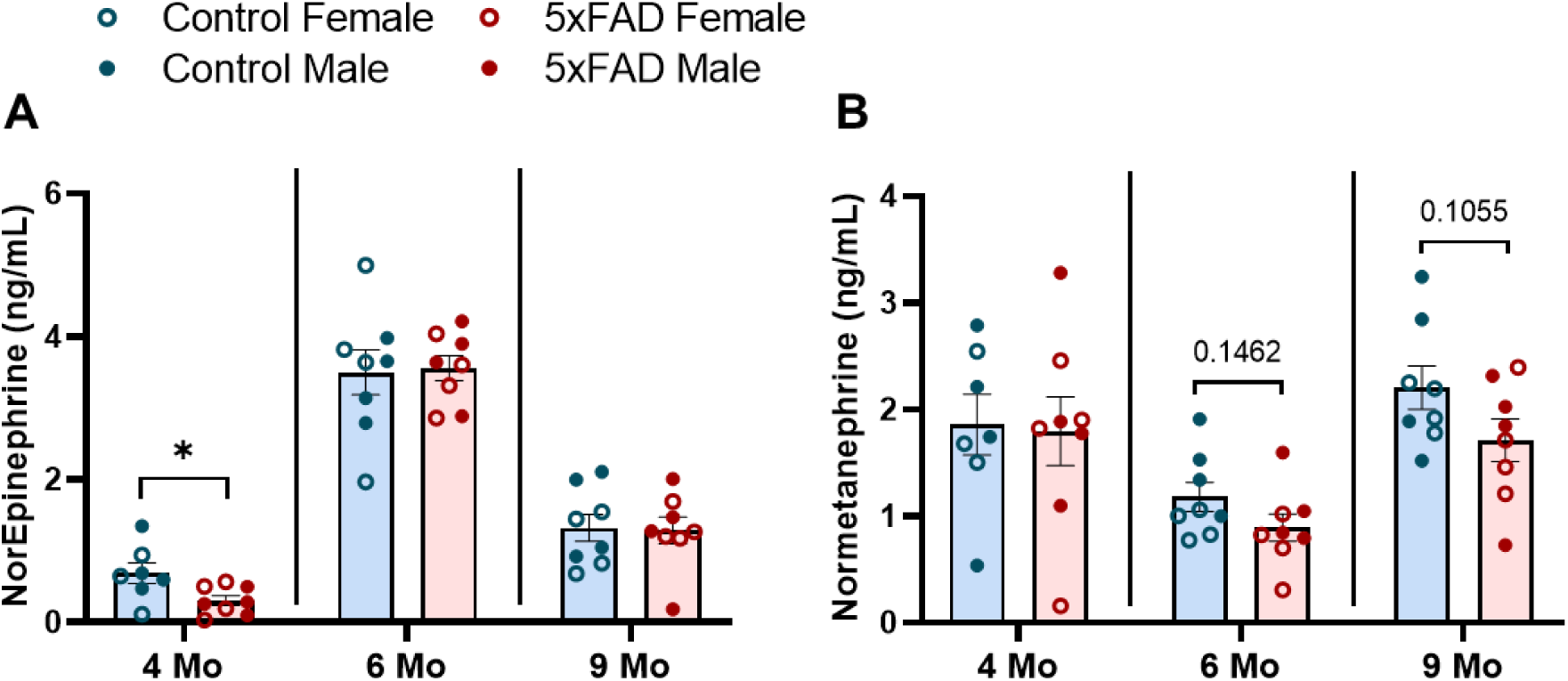
Reduced cortical NE and NMN levels in 5xfAD mice. Cortical lysates from all three age groups were used as substrates for ELISA for NE (**A**) and NMN (**B**). Cortices from different age groups were collected at different times, and thus, analyzed independently (separated by vertical lines). *n = 7-9, Student t-test; *p<0.05*.

**Supplementary Fig. 3:**
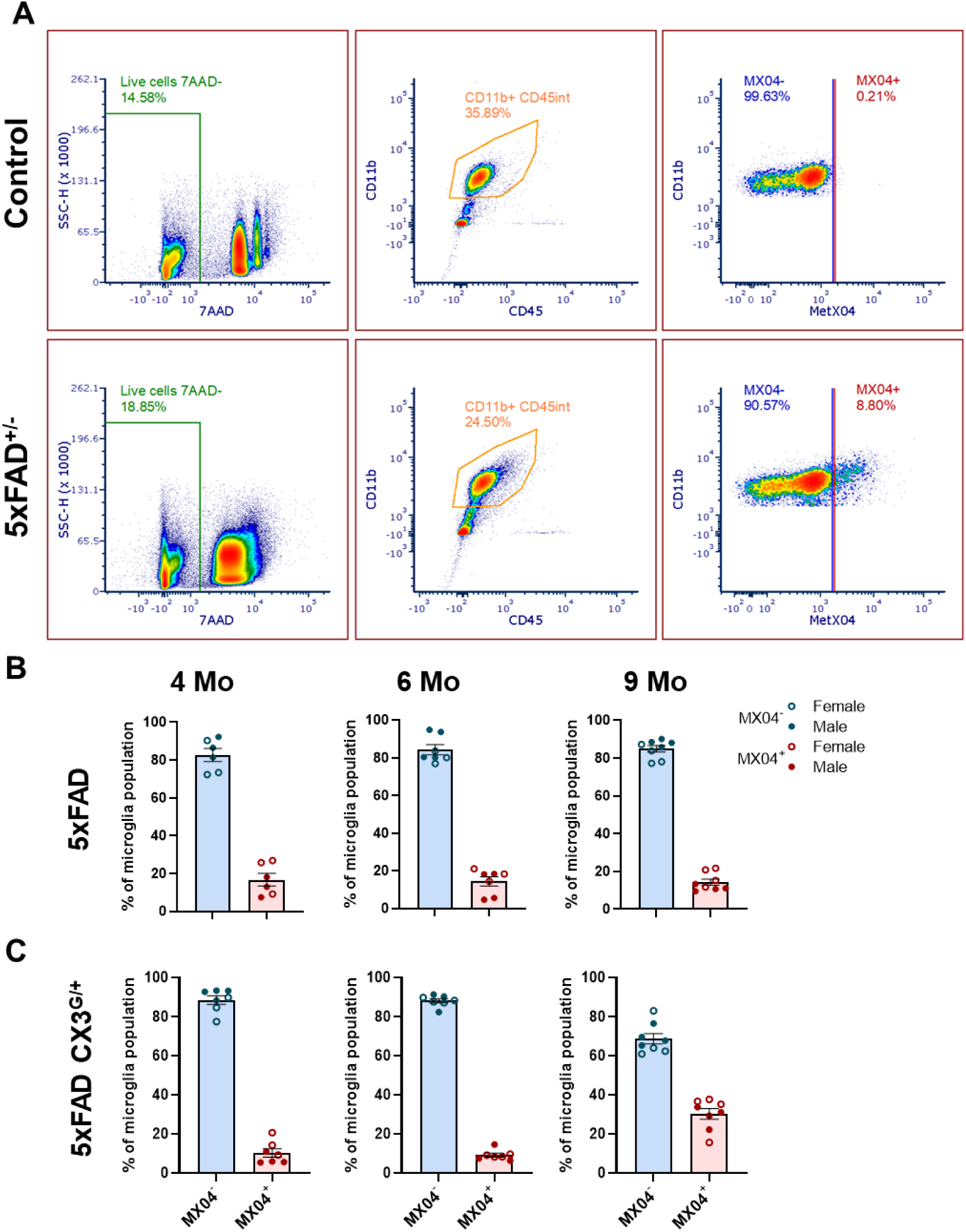
Gating strategy for microglia isolation with FACS in control (top panels) and 5xFAD (bottom panels) mice (**A**). Briefly, after debris exclusion, live cells were selected by gating for singlets and 7-AAD^−^ events. Microglia were defined as CD45^int^ CD11b^+^. MeX04 positive and negative events were determined with reference to 5xFAD, or non-transgenic C57BL/6 controls injected with MeX04 or vehicle (i.e., FMOs and negative controls). Representative samples from the 4-month-old cohort of 5xFAD and littermate control mice are shown here. All flow cytometry experiments were gated as described above. Fraction of MeX04^+^ and MeX04^-^ microglia in brains of 5xFAD (**B**) and 5xFAD CX3^G/+^ (**C**) mice across all age groups. *n = 7-9*.

**Supplementary Fig. 4:**
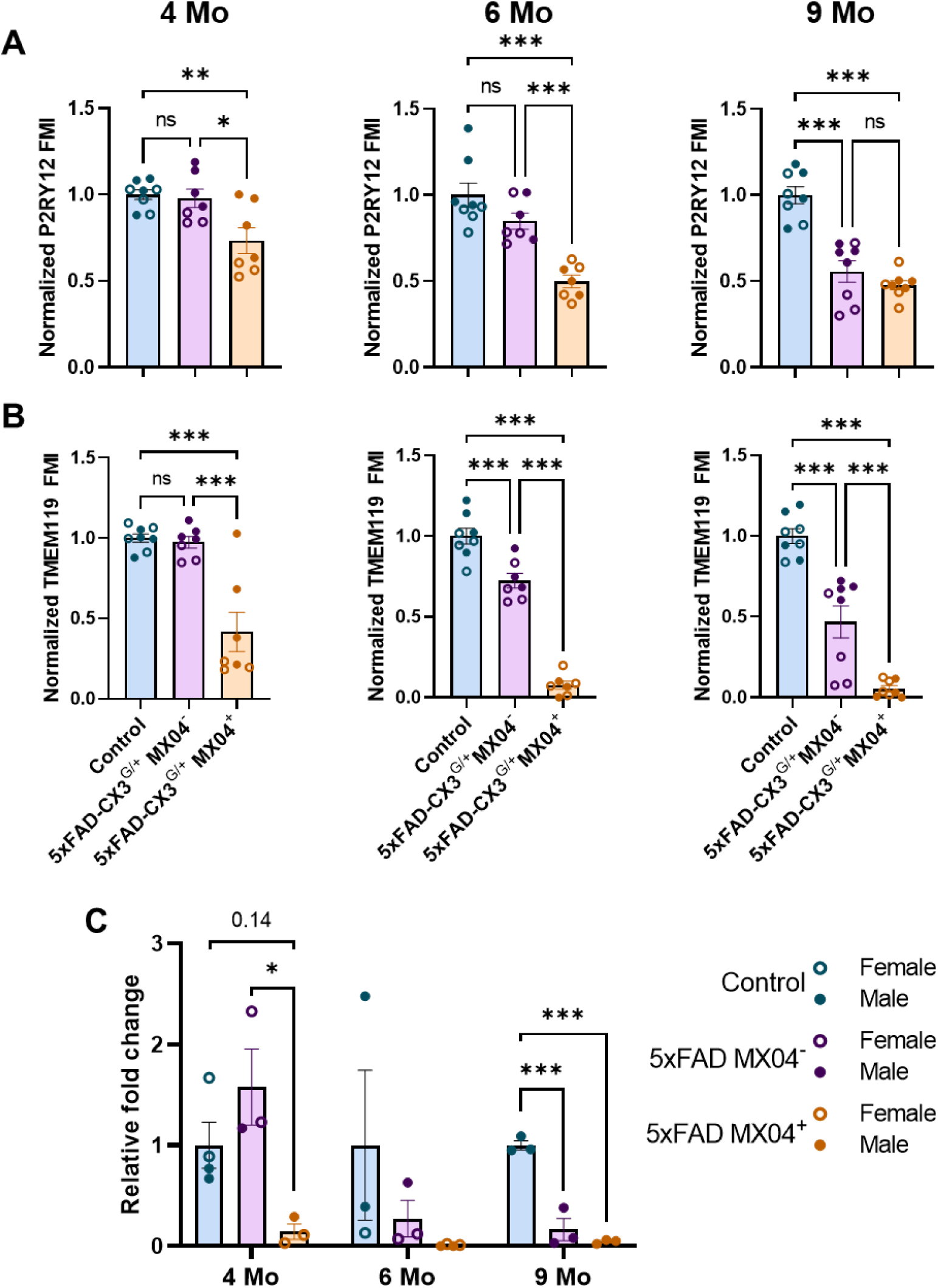
Amyloid pathology and age-dependent synergistic loss in microglial homeostatic signature and β2AR expression in 5xFAD CX3^GFP/+^ mice. Expression of microglial homeostatic marker P2RY12 (**A**) and TMEM119 (**B)**. Expression of β2AR in isolated microglia measured by qPCR (**C**). *n = 7-9 for FACS, n=3-4 for qPCR, one-way ANOVA with Bonferroni post-hoc correction, *p<0.05, **p<0.01, ***p<0.001*.

**Supplementary Fig. 5:**
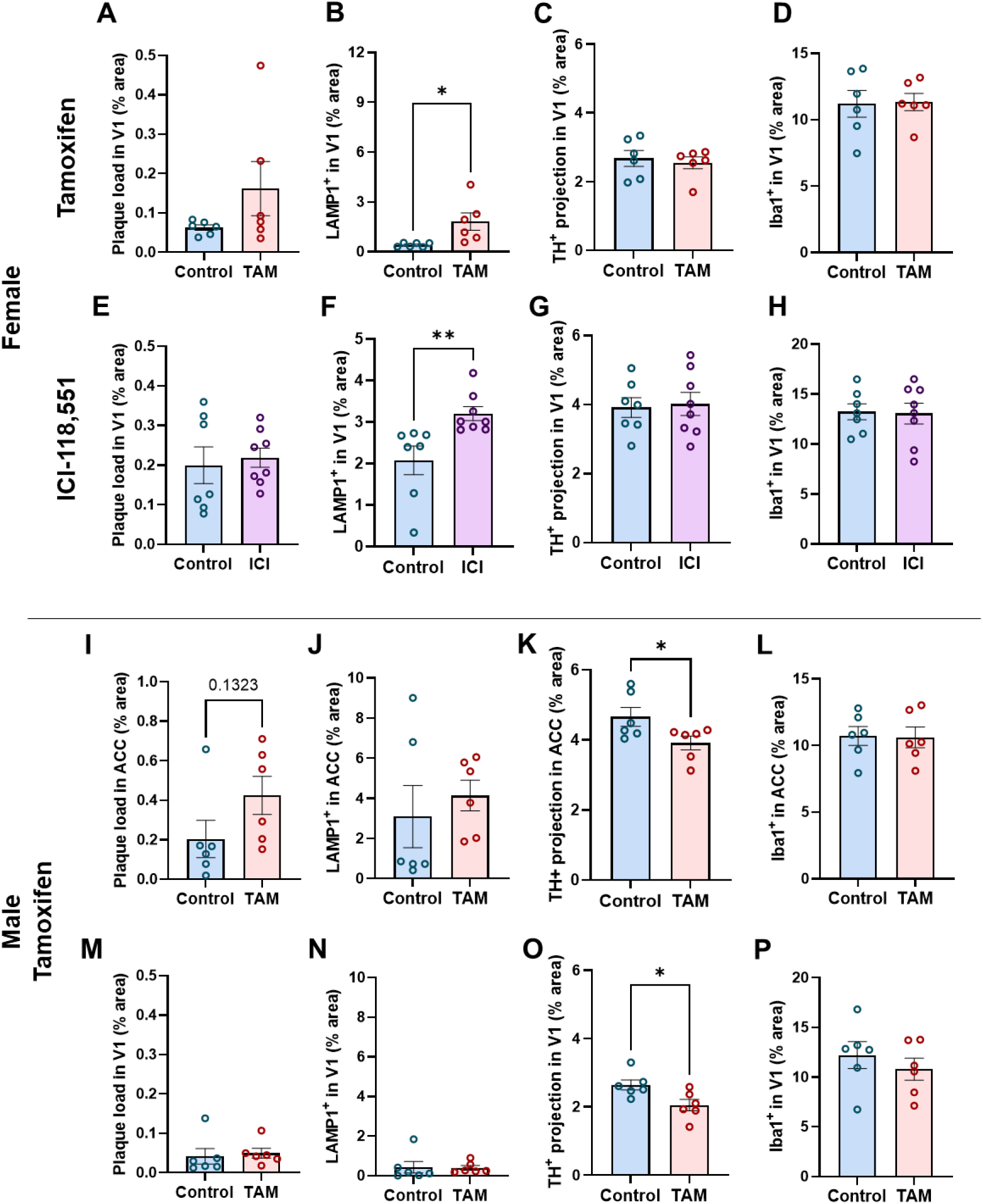
Inhibition of microglial β2AR signaling exacerbates amyloid pathology-associated damages differently between female and male 5xFAD mice. Quantification of plaque load, neuritic damage, TH^+^ projections and microglia activation in V1 of control and treated female 5xFAD animals, with microglia-specific deletion of β2AR (**A-D**) or prolonged 1-month treatment with ICI-118,551, a β2AR-specific antagonist (**E-H**). Quantification of plaque load, neuritic damage, TH^+^ projections and microglia activation in the ACC (**I-L**) and V1 (**M-P**) of control and treated male 5xFAD animals, with microglia-specific deletion of β2AR. *n = 6-8, Student t-test; *p<0.05, **p<0.01; Scale bar: 200μm*

**Supplementary Fig. 6:**
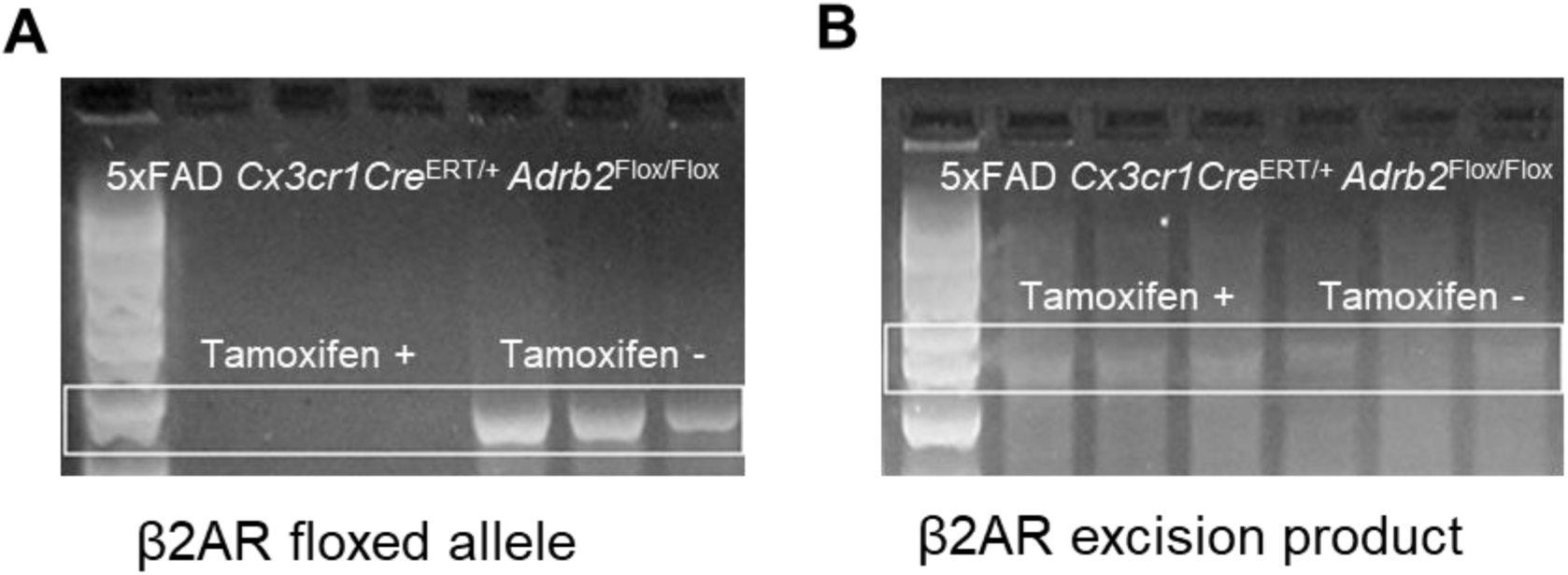
CX3CR1Cre^ERT^ successfully drives excision of β2ARs in microglia. PCR for the presence of the β2AR floxed allele. In mice given tamoxifen the allele is no longer present, suggesting excision (**A**). Recombination occurs in both the tamoxifen-treated and untreated mice due to leakiness of the Cre in this line (**B**). *n = 3 biological replicates*.

**Supplementary Fig. 7:**
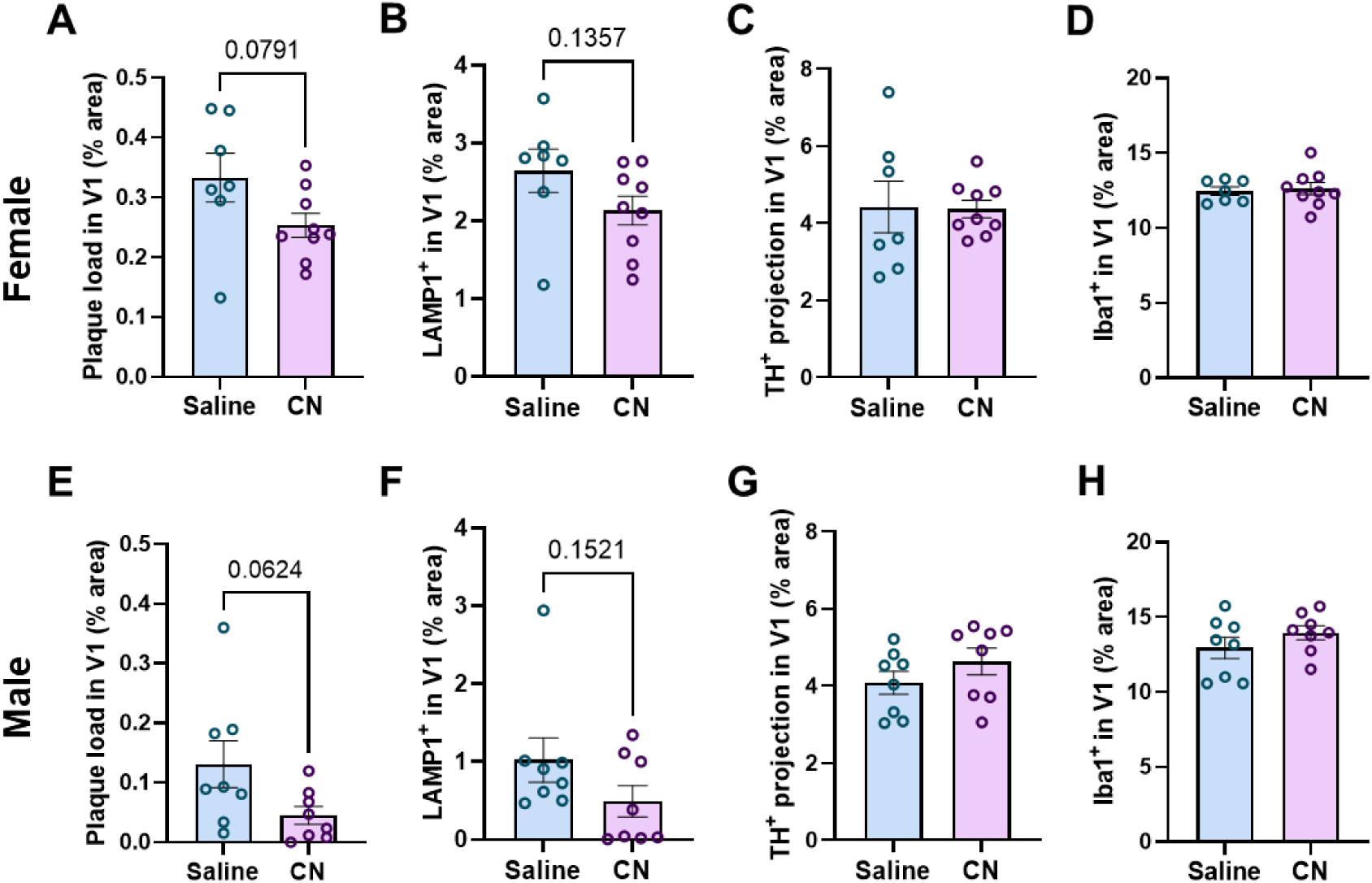
Prolonged exposure to β2AR agonist modestly attenuates amyloid pathology and associated neuritic damage in V1 in 5xFAD mice. Quantification of plaque load, neuritic damage, TH^+^ projection and microglia activation in cortex of control and treated 5xFAD females (**A-D**) and males (**E-H**) with prolonged 1-month treatment with Clenbuterol, a potent β2AR agonist. *n = 8-9, Student t-test; *p<0.05, **p<0.01; Scale bar: 200μm*

## Notes

### Competing Interest Statement

The authors have declared no competing interest.

